# Repertoire and abundance of secreted virulence factors shape the pathogenic capacity of *Pseudomonas syringae* pv. *aptata*

**DOI:** 10.1101/2023.03.20.533544

**Authors:** Ivan Nikolić, Timo Glatter, Tamara Ranković, Tanja Berić, Slaviša Stanković, Andreas Diepold

## Abstract

*Pseudomonas syringae* pv. *aptata* is a member of the sugar beet pathobiome and the causative agent of leaf spot disease. Like many pathogenic bacteria, *P. syringae* relies on the secretion of toxins, which manipulate host-pathogen interactions, to establish and maintain an infection. This study analyzes the secretome of six pathogenic *P. syringae* pv. *aptata* strains with different defined virulence capacities in order to identify common and strain-specific features, and correlate the secretome with disease outcome. All strains show a high type III secretion system (T3SS) and type VI secretion system (T6SS) activity under apoplast-like conditions mimicking the infection. Surprisingly, we found that low pathogenic strains show a higher secretion of most T3SS substrates, whereas a distinct subgroup of four effectors was exclusively secreted in medium and high pathogenic strains. Similarly, we detected two T6SS secretion patterns: while one set of proteins was highly secreted in all strains, another subset consisting of known T6SS substrates and previously uncharacterized proteins was exclusively secreted in medium and high virulence strains. Taken together, our data show that *P. syringae* pathogenicity is correlated with the repertoire and fine-tuning of effector secretion and indicate distinct strategies for establishing virulence of *P. syringae* pv. *aptata* in plants.

## INTRODUCTION

*Pseudomonas syringae* is the most ubiquitous bacterial plant pathogen on Earth (Mansfield et al., 2012). Numerous pathogenic varieties (pathovars) of *P. syringae* infect a plethora of different host plants worldwide. The life cycle of *P. syringae* is characterized by an epiphytic and an endophytic phase (Xin et al., 2018). The epiphytic phase encompasses bacterial survival and proliferation in fluctuating environmental conditions at the leaf surface and is characterized by the effort to gain access to the apoplast, an intercellular space filled with air and water, through leaf openings (stomata, hydathodes or wounds). By entering the apoplast, *P. syringae* starts the endophytic phase, establishing a parasitic lifestyle and proliferating in this nutrient-accessible environment (Baltrus et al., 2017).

Like many other Gram-negative bacterial plant pathogens (Büttner & He, 2009), *P. syringae* uses its type III secretion system (T3SS) for infection and suppression of plant immunity (Dillon et al., 2019). The T3SS is a syringe-like structure that injects virulence proteins effector proteins directly into the cytosol of eukaryotic target cells (Galán & Wolf-Watz, 2006; Büttner, 2012; Liu et al., 2016; Deng et al., 2017). The T3SS apparatus, also referred to as injectisome, is comprised of five main functional modules: (i) a cylindrical basal body that spans the bacterial inner and outer membranes (Marlovits et al., 2004) and encompasses (ii) an export apparatus in the inner membrane that likely acts as main secretion barrier (Hüsing et al., 2021), (iii) a dynamic cytosolic complex with six injectisome-bound pod substructures whose subunit exchange with cytosolic pools (Milne-Davies et al., 2021), (iv) a needle/pilus structure that extrudes from the bacterial outer membrane and functions as a conduit for effector transfer (Loquet et al., 2012), and (v) a translocon complex that produces a pore in the target cell plasma membrane to inject the effectors (Liu et al., 2016).

In *P. syringae*, the extracellular pilus is helically assembled from hundreds of copies of small HrpA proteins (around 11 kDa), which form a long narrow channel (2 μm in length, 6–8 nm in diameter) (Xin et al., 2018). While the HrpA sequence varies among pathovars of *P. syringae*, possibly as an adaptation to different cuticular and cell wall barriers in host plants, HrpA proteins in different pathovars share similar physicochemical and structural properties (Guttman et al., 2006). *P. syringae* strains have an exceptionally large variety of T3SS effector proteins, classified into 70 different families (Bundalovic-Torma et al., 2022), of which individual strains express 15-30 effectors (Wei et al., 2018). The four effectors encoded in the conserved Hrp^1^ pathogenicity island effector locus (AvrE, HopI, HopAA and HopM)^2^ are considered as core effectors, whereas the majority of effectors are encoded elsewhere in the genome (Lindeberg et al., 2012). Besides the virulence effectors, plant pathogens secrete T3SS helper proteins (harpins) involved in effector translocation into target cells. The diversity in the T3SS arsenal indicates that the strain-specific effector proteins repertoire plays an important role in host specificity and overall severity of *P. syringae* infections. However, effectors and harpins can also activate plant-triggered defense mechanisms (effector-triggered immunity, ETI) (Laflamme et al., 2020), such as the hypersensitive response (Choi et al., 2013). Effectors which could elicit ETI are widespread in *P. syringae* populations. A recent study showed that 97.2% of the 494 genomes of *P. syringae* strains organized in the *P. syringae* Type III Effector Compendium (PsyTEC) possess at least one orthologue of putative ETI-eliciting alleles, with 74.3% of strains encoding more than one (up to eight) putative ETI-elicitors (Martel et al., 2022).

In addition to the T3SS, type VI secretion systems (T6SS) have been attributed a major role in virulence and plant colonization of *P. syringae*; their importance in inter-bacterial competition has also recently been highlighted (Chien et al., 2020). Structurally, the T6SS resembles a phage tail-like device formed by a rigid tube of hexameric Hcp (TssD) rings wrapped in a contractile sheath consisting of TssB and TssC. Upon contraction of the sheath, the Hcp tube, which is sharpened by a tip of the VgrG (TssI) and PAAR proteins, is propelled outside the cell, and into a target cell (Cherrak et al., 2019). While the membrane complex is stable and can be reused for multiple injections, the contracted sheath is then disassembled by a ClpV (TssH) ATPase, and recycled for a new assembly and injection cycle (Zoued et al., 2014). Hcp and VgrG proteins are detectable in the extracellular medium of T6SS-active bacteria, which allows monitoring of T6SS activity *in vitro* (Pukatzki et al., 2006). Hcp and VgrG can carry additional C-terminal extension regions acting as effector domains (Pukatzki et al., 2007; Blondel et al., 2009). Such so-called specialized Hcp proteins with effector domains are common among Enterobacteriaceae (Ma et al., 2017), while specialized VgrG proteins are widespread in Betaproteobacteria and Gammaproteobacteria (Lien & Lai, 2017). Other T6SS effectors are so-called cargo effectors, which can be secreted by association with the exported T6SS components. Some cargo effectors, such as rearrangement hotspot (Rhs) and related YD-peptide repeat proteins, are often associated with an N-terminal PAAR domain or downstream of VgrG and PAAR-encoding genes (Koskiniemi et al., 2013; Pei et al., 2020). In contrast, most cargo effectors are encoded dispersed in the genome, usually next to a gene encoding their cognate immunity protein, which prevents self-harm. Accordingly, many T6SS effectors still await their identification. Bacteria, including plant pathogens such as *P. syringae, Agrobacterium tumefaciens*, and *Pantoea anantis*, can harbor multiple T6SS clusters in single strains (Sarris et al., 2010; Chien et al., 2020); however, other *P. syringae* pathovars contain only one T6SS cluster (Bernal et al., 2018).

*P. syringae* pv. *aptata* strains are causative agents of leaf spot disease, which affects several hosts, such as sugar beet, Swiss chard, cantaloupe and squash (Morris et al., 2000; Koike et al., 2003; Sedighian et al., 2014; Nikolić et al., 2018). Symptoms of the disease include circular or irregular necrotic spots on leaves, with a light-brown center and glassy black margins in the early stage (Stojšin et al., 2015). Mounting evidence for disease occurrence in sugar beets fields worldwide indicates the robust pathogenic potential of this particular *P. syringae* pathovar (Stojšin et al., 2015; Arabiat et al., 2016; Rotondo et al., 2019; Nampijja et al., 2021). A previous comparative characterization of individual strains from a collection of *P. syringae* pv. *aptata* found high diversity in terms of pathogenicity and host range (Nikolić et al., 2018; Morris et al., 2019). In particular, pathogenicity assays and measurements of disease severity conducted in greenhouse experiments revealed distinctive groups of strains, classified as low, intermediate and high virulence. The high virulence strains display a broad host range (10-16 plant species), whereas low virulence strains caused disease in one or few tested host plants, indicating a narrow host range (Nikolić et al., 2018; Morris et al., 2019). In this report, we analyze the secretome of six selected *P. syringae* pv. *aptata* strains covering a wide range of distinctive pathogenic characteristics (low/high virulence and narrow/broad host range) under *in vitro* conditions that mimic the apoplast environment, in order to link the repertoire of secreted effector proteins and the disease outcome. To this aim, we performed the first in-depth secretome analysis of the emerging sugar beet pathogen *Pseudomonas syringae* pv. *aptata* and focused on the identification and quantification of T3SS and T6SS effectors and motility factors. Notably, we found that the investigated low virulence strains of *P. syringae* pv. *aptata* secreted large amounts of a defined spectrum of type III secretion effectors, whereas high virulence strains have a broader spectrum of effectors generally secreted in lower abundance. While all strains showed high T6SS activity, medium and high virulence strains secreted an additional subset of known T6SS effectors and previously uncharacterized co-secreted proteins that are candidates for novel *P. syringae* T6SS effectors. This identification of distinct protein secretion profiles may reveal defined infection strategies, which correlate with the infection severity and host range of *P. syringae* pv. *aptata*.

## MATERIAL AND METHODS

### Strain collection and growth conditions

The six *P. syringae* pv. *aptata* strains used in this work display different pathogenic capacities regarding virulence and host range. Strains P16 and P17 show low virulence properties (10% of leaf plate under necrotic symptoms 7 days after inoculation) with a narrow host range (disease caused in 1 and 2 out of 16 tested plant species), while strains P21 and P23 represent highly virulent strains (70% of necrotic leaf plate) with broad host range (disease caused in all 16 plant species); strains P26 and P93 showed intermediate values for virulence and host range (35% of necrotic leaf plate; disease caused in 5 of the 16 plant species) (Nikolić et al., 2018; Morris et al., 2019). T3SS secreting conditions mimicking the apoplast-like environment were established by growing cells in *hrp*-inductive minimal medium (HIM) containing 10 mM fructose at pH 5.7 at 22°C, as defined in (Huynh et al., 1989). King’s B medium (KB) (King et al., 1954) at 28°C and neutral pH was used for non-secreting conditions. For the analysis of the secretome in secreting and non-secreting conditions, three biological replicates of 50 ml cultures (OD_600_= 0.15) of each strain were grown on a shaking incubator (180 rpm) in both conditions for six hours, while two biological replicates grown in the same conditions were used for the secretome comparisons between tested strains.

### Proteomics analysis

50 ml of culture grown as described above were collected by centrifugation at 4500 x g for 10 min at room temperature. Supernatants were transferred into fresh 50 ml tubes and filtered through 0.22 µm filters twice. Proteins for the secretome analysis were precipitated by the addition of 10% (v/v) trichloroacetic acid (TCA) and incubation at 4°C overnight. Precipitated proteins were collected by centrifugation for 30 min at 4000 x g and 4°C; the protein pellet was washed twice with 5 ml ice-cold acetone. Dried protein pellets were solubilized in 500 μl reconstitution buffer (1 M Urea in 100 mM ammonium bicarbonate (ABC) + 5 mM Tris(2-carboxyethyl) phosphine (TCEP)), and sonicated for 20 s (Hielscher Ultrasound Technology). Protein reduction by the added TCEP was carried out for 30 min at 37°C, followed by alkylation using iodoacetamide (10 mM) for 30 min at 25°C in the dark. For protein digestion, 1 μg trypsin (Sequencing grade Modified Trypsin, Promega) was added to the sample, and incubated at 30°C overnight. After the digest, samples were acidified using 1.5% (v/v) trifluoroacetic acid (TFA) and C18-purified using Micro Spin Columns (Harvard Apparatus) according to the manufacturer’s instructions.

The protein database for the proteomics analysis is based on the protein list for strain *P. syringae* pv. *aptata* ICMP459 (UniProt identifier UP000050439), also isolated from sugar beet. Protein sequences of T3SS effectors from the proteomes of the other four *P. syringae* pv. *aptata* proteome datasets available at the UniProt database (strain G733/Uniprot identifier UP000271836, ICMP11935/UP000274315, ICMP4388/UP000274541 and DSM50252/UP000005484) and effectors from other *P. syringae* pathovars, obtained from the publicly available *P. syringae* effector database^3^ were added to this list. The final protein list (Suppl. File 1) contains 5243 proteins, including 63 T3SS effectors.

Analysis of raw data for label-free quantification analysis was performed using MaxQuant (Cox & Mann, 2008) in standard settings using the assembled protein database. Carbamidomethylation (C) was set as fixed, oxidation (M) and deamidation (N, Q) as variable modifications. The proteomics data have been deposited to the ProteomeXchange Consortium via the PRIDE partner repository with the dataset identifier PXD040715.

For spectral based assessment searches were carried out using Mascot (v2.5, Matrix Science) with 10 ppm MS1 and 0.02 Da fragment ion tolerance with Carbamidomethylation (C) as fixed, oxidation (M) and deamidation (N, Q) as variable modifications. Search results were evaluated in Scaffold 4 (Proteome Software).

### T3SS pili purification

150 ml day cultures of strain *P. syringae* pv. *aptata* P17 in T3SS were grown in secreting and non-secreting conditions. The cultures were transferred to 50 ml conical tubes and cells were collected by centrifugation at 3000 x g for 8 min at 4°C. Pellets were vigorously resuspended in 800 µl Tris-HCl buffer (20 mM, pH 7.5), leading to shearing of the pili, analogous to the purification of T3SS needles in *Yersinia enterocolitica*. The resuspension was transferred to 2 ml Eppendorf tubes and centrifuged at 4000 x g for 6 min at 4°C. The supernatant containing the pili was transferred to 1.5 ml Eppendorf tubes and centrifuged at 24,100 x g for 60 min at 4°C. The pellet was resuspended in 65 µl 1 x SDS-protein loading buffer (Laemmli, 1970). Prepared samples were stored at -20°C prior to analysis.

### SDS-PAGE analysis

15 % resolving gels (5 ml 30 % acrylamide; 2.16 ml dH_2_O; 2.5 ml 1.5 M Tris-HCl buffer pH 8.8; 400 µl 10 % SDS; 75 µl 10 % APS; 40 µl TEMED) were prepared, poured into a casting chamber and overlaid with 1 ml of isopropyl alcohol to ensure a uniform gel boundary. The gels were washed with distilled water after 30 min polymerization to remove isopropyl alcohol and un-polymerized acrylamide. The stacking gel (1.16 ml 30 % acrylamide; 2.56 ml dH_2_O; 1.16 ml 0.5 M Tris-Cl buffer pH 6.8; 600 µl 10 % SDS; 25 µl 10 % APS; 20 µl TEMED) was poured over the resolving gel.

### Electron microscopy

10 ml day cultures of *P. syringae* pv. *aptata* P17 were grown in secreting and non-secreting conditions. Formvar coated copper grids (Plano, S162-3, 300 mesh) were hydrophilized using a PELCO easiGlow™ Glow Discharge Cleaning System. 5 µl cell suspension was applied to the grid and incubated for 1 min, before the excess liquid was removed with Whatman paper. The grids were negatively stained using 2% uranyl acetate and washed once with ddH_2_O. Cells were imaged using a JEM-1400 electron microscope (JEOL, Japan) at 100 kV. Determination of the length and diameter of flagella and T3SS pili was performed by ImageJ software (Schneider et al., 2012).

### Swimming and swarming assays

Bacterial cultures were grown overnight (30°C with shaking at 180 rpm) in KB and diluted in fresh medium to adjust OD_600_ ∼0.3 (approximately 2 × 10^8^ colony forming units/ml) for motility assays. The preparation of swimming and swarming media and the experimental procedure were performed as previously described by (Hockett et al., 2013). In brief, swimming media was prepared as 50% KB containing 0.25% agar; swarming media was prepared as undiluted KB containing 0.4% agar. Swimming assays were performed by stabbing bacterial cell suspension in the center of a swimming plate with a sterile 10 µl pipette tip. For the swarming assay, 3 µl aliquots were inoculated onto the center of a swarming plate. Each experiment was done in triplicates. Swimming and swarming motilities were observed 24 h after incubation at room temperature. Swimming diameters were measured using ImageJ software (Schneider et al., 2012).

### Bioinformatics Analysis

To identify putative T6SS effectors, the whole *P. syringae* pv. *aptata* proteome of strain P21, excluding proteins smaller than 50 amino acids and larger than 5000 amino acids, was analyzed with the Bastion6 pipeline (Wang et al., 2018). Proteins with a Bastion6 score ≥ 0.7 were considered as putative T6SS effectors. Genome mapping was performed by CGview web server (Grant & Stothard, 2008) using a *P. syringae* pv. *aptata* P21 genome assembly file (https://www.ncbi.nlm.nih.gov/assembly/GCF_018530765.1), while operon prediction was performed with the Operon-Mapper web server (Taboada et al., 2018) using the P21 genome sequence.

## RESULTS

### *P. syringae* pv. *aptata* forms T3SS pili and secretes T3SS effectors under apoplast-like conditions

To test the requirements for the formation of T3SS HrpA pili and the secretion of potential effectors, we incubated the low virulence strain *P. syringae* pv. *aptata* P17 under apoplast-like conditions activating T3SS secretion (Huynh et al., 1989) (secreting conditions) and non-secreting conditions. Electron microscopy analysis showed the formation of pili (7.3 ± 0.8 nm in diameter) exclusively under secreting conditions (Fig. 1A, Suppl. Table 1). Analysis of the culture supernatant of bacteria grown under T3SS secreting conditions on SDS-PAGE gels identified bands that were exclusively visible in the supernatant of bacteria grown under secreting conditions, including a band running at a molecular weight of approximately 15 kDa, compatible with the size of HrpA (expected molecular weight of 11 kDa) and additional weaker bands at higher molecular weights (Fig. 1B). To reveal the identity of these proteins and to get an overview of the secretome of *P. syringae* pv. *aptata*, we carried out a shotgun proteomics mass spectrometry approach. The results confirmed the specific secretion of HrpA under T3SS secreting conditions. Importantly, they further revealed the specific secretion of several additional known T3SS effectors (Table 1). Similar to HrpA, the majority of these effectors (HopBA1, HopM1, HopBF1, HopI1, HopAA1, HopC1, HopAZ1, HopZ3, HopPmaK, HopBC1) were only detected under secreting conditions. One effector (HopJ1) was detected in low amounts under both conditions, while one (HopAH2) was detected in higher amounts under non-secreting conditions.

**Figure 1.**
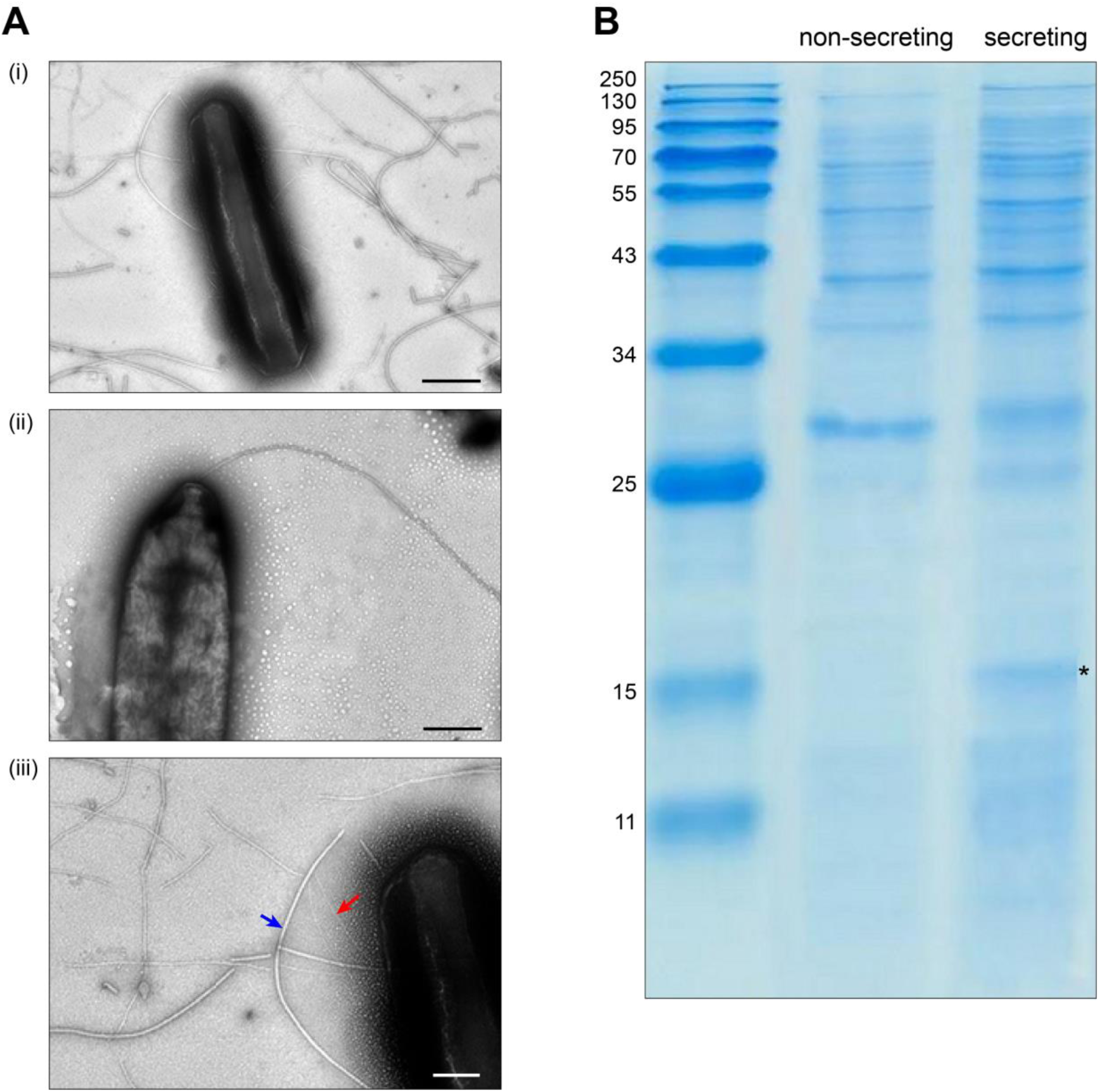
Specific T3SS protein secretion and pilus formation under apoplast-like conditions. **(A)** Transmission electron microscopy of *P. syringae* pv. *aptata* P17 after incubation (i) in secreting conditions, (ii) in non-secreting condition (scale bars, 500 nm). (iii) Magnification of (i) showing the presence of T3SS pili (red arrow) with an observed diameter of 7.3 ± 0.8 nm, in comparison to flagella (blue arrow) with an observed diameter of 18.0 ± 4.0 nm (scale bar, 200 nm). (**B**) SDS-PAGE analysis of culture supernatant of *P. syringae* pv. aptata P17. Left, molecular weight marker (kDa); asterisk denotes additional band compatible with the molecular weight of HrpA (11.0 kDa).

**Table 1:**
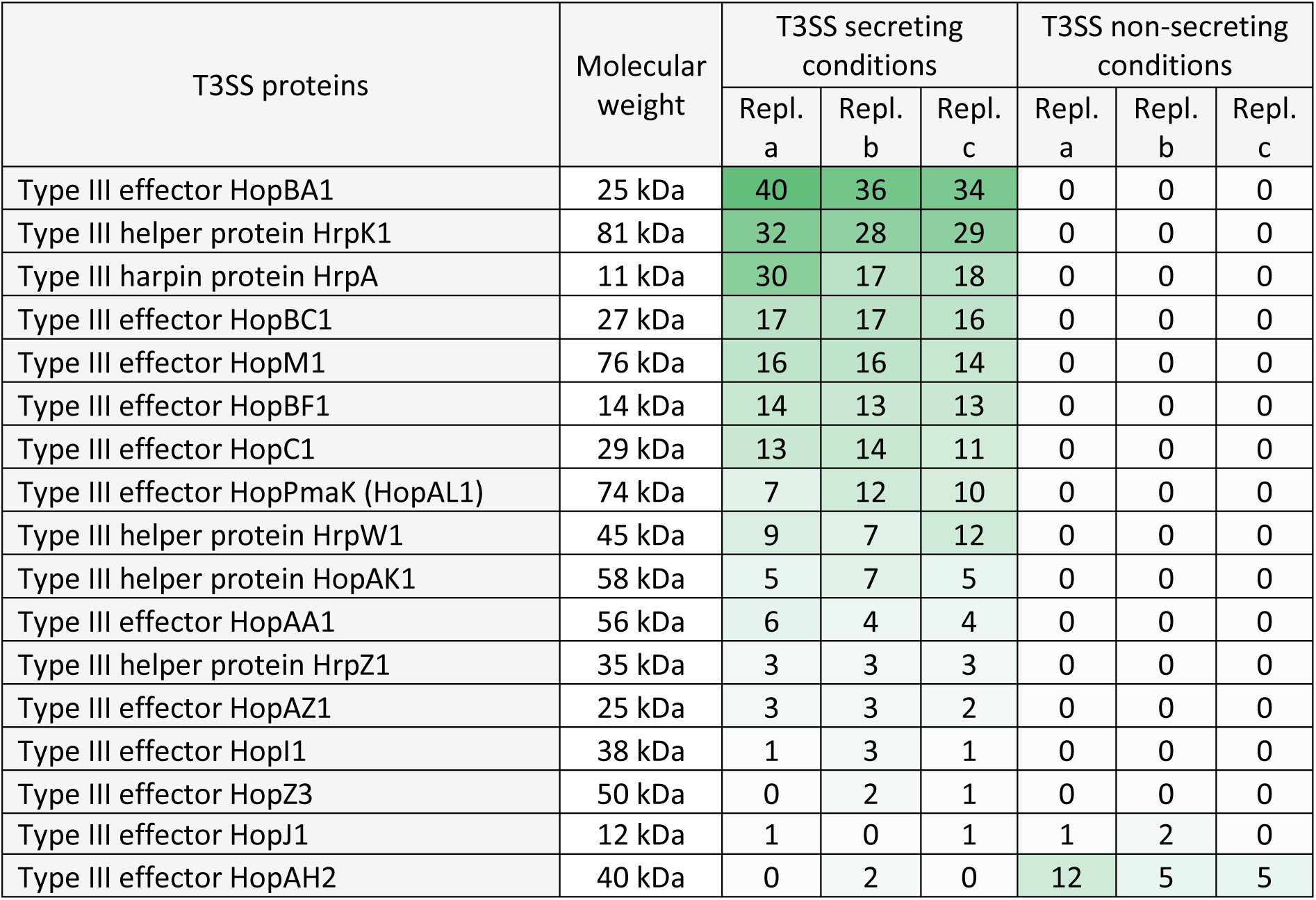
Detection of T3SS in culture supernatant under different conditions. LC-MS/MS-based quantification of biological triplicates (replicates a-c) of *P. syringae* pv. *aptata* strain P17. T3SS effector peptide counts detected in the culture supernatant in secreting or non-secreting conditions, as indicated.

| T3SS proteins | Molecular weight | T3SS secreting conditions |  |  | T3SS non-secreting conditions |  |  |
| --- | --- | --- | --- | --- | --- | --- | --- |
|  |  | Repl. a | Repl. b | Repl. c | Repl. a | Repl. b | Repl. c |
| Type III effector HopBA1 | 25 kDa | 40 | 36 | 34 | 0 | 0 | 0 |
| Type III helper protein HrpK1 | 81 kDa | 32 | 28 | 29 | 0 | 0 | 0 |
| Type III harpin protein HrpA | 11 kDa | 30 | 17 | 18 | 0 | 0 | 0 |
| Type III effector HopBC1 | 27 kDa | 17 | 17 | 16 | 0 | 0 | 0 |
| Type III effector HopM1 | 76 kDa | 16 | 16 | 14 | 0 | 0 | 0 |
| Type III effector HopBF1 | 14 kDa | 14 | 13 | 13 | 0 | 0 | 0 |
| Type III effector HopC1 | 29 kDa | 13 | 14 | 11 | 0 | 0 | 0 |
| Type III effector HopPmaK (HopAL1) | 74 kDa | 7 | 12 | 10 | 0 | 0 | 0 |
| Type III helper protein HrpW1 | 45 kDa | 9 | 7 | 12 | 0 | 0 | 0 |
| Type III helper protein HopAK1 | 58 kDa | 5 | 7 | 5 | 0 | 0 | 0 |
| Type III effector HopAA1 | 56 kDa | 6 | 4 | 4 | 0 | 0 | 0 |
| Type III helper protein HrpZ1 | 35 kDa | 3 | 3 | 3 | 0 | 0 | 0 |
| Type III effector HopAZ1 | 25 kDa | 3 | 3 | 2 | 0 | 0 | 0 |
| Type III effector HopI1 | 38 kDa | 1 | 3 | 1 | 0 | 0 | 0 |
| Type III effector HopZ3 | 50 kDa | 0 | 2 | 1 | 0 | 0 | 0 |
| Type III effector HopJ1 | 12 kDa | 1 | 0 | 1 | 1 | 2 | 0 |
| Type III effector HopAH2 | 40 kDa | 0 | 2 | 0 | 12 | 5 | 5 |

These results validate the induction of secretion by the apoplast-like conditions used in our experiments and allow an evaluation of our hypothesis that different pathogenic traits are correlated with the secretion of different virulence factors.

### Analysis of the secretome of *P. syringae* pv. *aptata* strains with different virulence

Based on the results described above, we reasoned that differences in the pathology of different strains of *P. syringae* pv. *aptata* might be correlated with the number, nature and/or amount of translocated effector proteins. To reveal such a correlation, we performed a label-free quantitative (LFQ) proteomics analysis of the secretome in apoplast-like conditions of six *P. syringae* pv. *aptata* strains with different pathogenic properties previously established (Nikolić et al., 2018; Morris et al., 2019) – the low virulence strains P16 and P17, medium virulence strains P26 and P93, and the high virulence strains P21 and P23. To focus on *bona fide* secreted proteins that might contribute to these differences, from an initial list of 901 detected proteins, we selected all proteins with at least five detected peptides and a maximal log_2_ intensity difference between individual strains of ≥3. The resulting list of 246 proteins largely excludes cytosolic proteins present at constant lower amounts due to limited cell lysis and includes the majority of known secreted proteins, such as 23 of 26 T3SS-secreted proteins, all six known T6SS-secreted proteins, and 8 of 15 exported flagellar proteins. These overall results confirm a significant and highly strain-specific contribution of known secretion systems to the secretome.

Flagellar proteins such as flagellin, hook-associated proteins and secreted regulatory components were found in the secretome of all strains in a more uniform quantity (Suppl. Table 2), with the highest levels secreted by the medium and high virulence strains (P23 and P93 > P21 and P26) and the lowest level by the low virulence strain P16 (Table 2). These results were partially reflected in swimming assays, where P16 had the lowest diameter, whereas P21 had the largest swimming diameter (Suppl. Fig. 1).

**Table 2:** Relative quantification of secreted proteins reveals specific secretion patterns. Shotgun LC-MS/MS and label-free quantification of proteins secreted by the T3SS, T6SS and flagella in the supernatant of the indicated *P. syringae pv. aptata* strains in secreting conditions. Proteins ordered by category and descending average detection intensity. Color scale indicates average protein intensities. White font indicates imputed values in the absence of detection. Different shades of green (for T3SS secreted proteins) or yellow (for T6SS secreted proteins) in the left column indicate different putative regulatory groups, see main text for details. LV, low virulence; MV, medium virulence; HV, high virulence.

| Protein | Average protein intensity (log 2)<br>in indicated strains |  |  |  |  |  | Overall<br>avg.<br>log 2<br>intens. | Max.<br>diff.<br>log 2<br>intens. |
| --- | --- | --- | --- | --- | --- | --- | --- | --- |
|  | P16<br>(LV) | P17<br>(LV) | P21<br>(HV) | P23<br>(HV) | P26<br>(MV) | P93<br>(MV) |  |  |
| Type III secretion system |  |  |  |  |  |  |  |  |
| Type III harpin protein HrpA | 35.158 | 35.424 | 33.986 | 31.331 | 32.991 | 30.108 | 33.166 | 5.317 |
| Type III helper protein HrpK1 | 33.692 | 32.764 | 31.349 | 28.746 | 29.748 | 28.417 | 30.786 | 5.276 |
| Type III helper protein HrpW1 | 33.116 | 31.356 | 28.468 | 28.810 | 28.660 | 28.344 | 29.792 | 4.772 |
| Type III helper protein HrpZ1 | 31.748 | 30.397 | 28.822 | 28.028 | 28.579 | 26.993 | 29.094 | 4.755 |
| Type III effector HopBA1 | 32.721 | 31.934 | 26.326 | 28.058 | 24.955 | 27.470 | 28.577 | 7.765 |
| Type III effector HopI1 | 30.611 | 29.790 | 28.913 | 26.368 | 28.069 | 25.733 | 28.247 | 4.878 |
| Type III effector HopBN1 | 31.312 | 30.115 | 28.620 | 25.265 | 27.861 | 25.648 | 28.137 | 6.047 |
| Type III effector AvrE1 | 29.827 | 28.419 | 28.164 | 26.723 | 27.696 | 25.853 | 27.780 | 3.973 |
| Type III helper protein HopAK1 | 31.575 | 30.297 | 27.908 | 25.085 | 26.734 | 24.926 | 27.754 | 6.649 |
| Type III effector HopAZ1 | 30.774 | 30.893 | 27.588 | 24.890 | 26.627 | 24.511 | 27.547 | 6.382 |
| Type III effector HopAA1 | 29.924 | 28.046 | 28.234 | 25.592 | 26.266 | 24.456 | 27.086 | 5.468 |
| Type III effector HopM1 | 30.460 | 29.606 | 26.765 | 25.864 | 25.604 | 23.930 | 27.038 | 6.530 |
| Type III effector HopBF1 | 32.206 | 30.950 | 25.168 | 26.384 | 21.845 | 25.634 | 27.031 | 10.361 |
| Type III effector HopC1 | 30.485 | 29.604 | 27.516 | 25.332 | 26.351 | 22.785 | 27.012 | 7.700 |
| Type III effector HopH1 | 29.731 | 28.448 | 25.761 | 24.816 | 24.829 | 23.583 | 26.195 | 6.148 |
| Type III effector HopBC1 | 28.918 | 28.708 | 25.921 | 24.103 | 24.917 | 24.094 | 26.110 | 4.825 |
| Type III inner rod protein HrpB | 27.842 | 26.900 | 26.291 | 23.718 | 24.533 | 22.243 | 25.254 | 5.599 |
| Type III effector HopZ3 | 27.853 | 28.582 | 22.023 | 25.252 | 21.296 | 25.160 | 25.028 | 7.286 |
| Type III effector HopAU1 (pv. theae) | 21.125 | 21.429 | 28.602 | 24.773 | 27.282 | 22.675 | 24.314 | 7.478 |
| Type III effector HopPmaK | 28.240 | 28.850 | 20.096 | 20.456 | 22.750 | 23.100 | 23.915 | 8.754 |
| Type III effector HopAW1 (pv. syringae) | 22.344 | 19.953 | 27.682 | 20.383 | 27.057 | 21.593 | 23.169 | 7.730 |
| Type III effector HopAH1 | 19.976 | 21.827 | 25.483 | 22.406 | 24.362 | 22.368 | 22.737 | 5.506 |
| Type III effector AvrRpm1 (pv. syringae) | 19.988 | 21.429 | 26.473 | 21.459 | 25.504 | 20.608 | 22.577 | 6.485 |
| Type VI secretion system |  |  |  |  |  |  |  |  |
| Putative type VI effector, Hcp1 family | 36.810 | 34.085 | 33.449 | 36.644 | 33.152 | 36.473 | 35.102 | 3.658 |
| Putative type VI effector, VgrG family | 30.412 | 29.552 | 28.495 | 30.660 | 27.655 | 30.453 | 29.538 | 3.005 |
| Putative type VI effector, Hcp1 family | 22.551 | 21.565 | 28.484 | 31.146 | 26.868 | 30.847 | 26.910 | 9.581 |
| Putative type VI effector, VgrG family | 23.417 | 23.397 | 20.542 | 27.142 | 22.382 | 26.237 | 23.853 | 6.600 |
| Rhs protein | 21.191 | 19.702 | 21.223 | 25.796 | 21.238 | 25.048 | 22.366 | 6.094 |
| Flagellum |  |  |  |  |  |  |  |  |
| Flagellar hook-associated protein 2 | 31.746 | 33.611 | 34.777 | 34.798 | 34.304 | 34.607 | 33.974 | 3.052 |
| Flagellar biosynthesis anti-sigma factor FlgM | 31.399 | 32.786 | 34.528 | 33.849 | 33.952 | 33.737 | 33.375 | 3.130 |
| Flagellar hook-associated protein FlgL | 28.693 | 31.349 | 30.997 | 32.219 | 31.066 | 31.774 | 31.016 | 3.526 |
| Flagellar hook-associated protein FlgK | 28.016 | 30.338 | 30.414 | 31.628 | 30.902 | 31.840 | 30.523 | 3.824 |
| Basal-body rod modification protein FlgD | 24.841 | 28.208 | 28.824 | 30.350 | 29.266 | 30.399 | 28.648 | 5.558 |
| Flagellar protein FlaG | 26.669 | 28.226 | 28.608 | 29.836 | 28.362 | 29.424 | 28.521 | 3.167 |
| Flagellar basal body protein | 23.260 | 25.782 | 25.413 | 27.150 | 25.913 | 27.403 | 25.820 | 4.144 |
| Flagellar rod assemb. prot./muramidase FlgJ | 22.328 | 22.479 | 25.790 | 25.060 | 24.430 | 24.772 | 24.143 | 3.462 |

Focusing on proteins secreted by the T3SS, we detected the HrpA pilus protein (which was the most intense T3SS protein in all strains), four abundant T3SS helper proteins (HrpK1, HrpW1, HrpZ1, and HopAK1), the regulatory component HrpB, and 17 different T3SS effector proteins (Table 2). The fact that all tested strains secreted core effectors from the conserved effector locus, such as AvrE1, HopM1, HopI1, and HopAA1, confirms the conserved presence of the complete *hrp* pathogenicity island encoding the T3SS. The majority of T3SS effectors were secreted in greatest relative amounts by the low pathogenicity strains P16 and P17, and to a lower degree (on average by a factor of 10-20 lower) by the medium and high virulence strains P21, P23, P26 and P93. However, a distinct subset of four effectors, HopAU1, HopAW1, HopAH1 and AvrRpm1, showed a clearly different pattern – these proteins were specifically detected in the supernatant of the medium and high pathogenicity strains P21 and P26 (Table 2).

For the T6SS, we detected large amounts of the secreted tube protein Hcp1 (which was, in fact, the protein with the highest average intensity across all strains) and the VgrG tip protein. Both proteins were most abundant in the secretome of strains P16, P23 and P93. In addition, another set of T6SS genes was secreted with a different pattern: Additional variants of Hcp1, VgrG, and an Rhs protein were all most abundant in strains P23 and P93 (Table 2). To better understand these results, we performed a bioinformatic analysis of T6SS genes in the known genome sequences of the strains used in this study. In the genome sequence of strain P21, T6SS genes are organized in two genomic islands (clusters) (Fig. 2), suggesting the presence of two types of T6SSs as already identified in other plant-associated bacteria (Bernal et al., 2018). The T6SS cluster 1 contains a single operon containing genes *tssA-tssM*, as well as *hcp* and *tagF*, encoding a T6SS accessory protein (TagF) involved in repression of T6SS in bacteria (Lin et al., 2018). The T6SS cluster 2 contains the operon from *tssA*-*tssM*, although differently arranged than in T6SS cluster 1, and a second operon with the *hcp* and *vgrG* genes. The Hcp1 and VgrG proteins detected with high intensities in the supernatant of all strains (Table 2) are highly homologous to T6SS cluster 1 *hcp* and *vgrG*, whereas the Hcp1 and VgrG variants detected in lower intensities and not across all strains cannot be unambiguously assigned to specific P21 genes. Other T6SS-related genes that encode putative T6SS-secreted proteins (Rhs, VgrG, Hcp) are encoded separately from T6SS clusters in P21 (Fig. 2).

**Figure 2.**
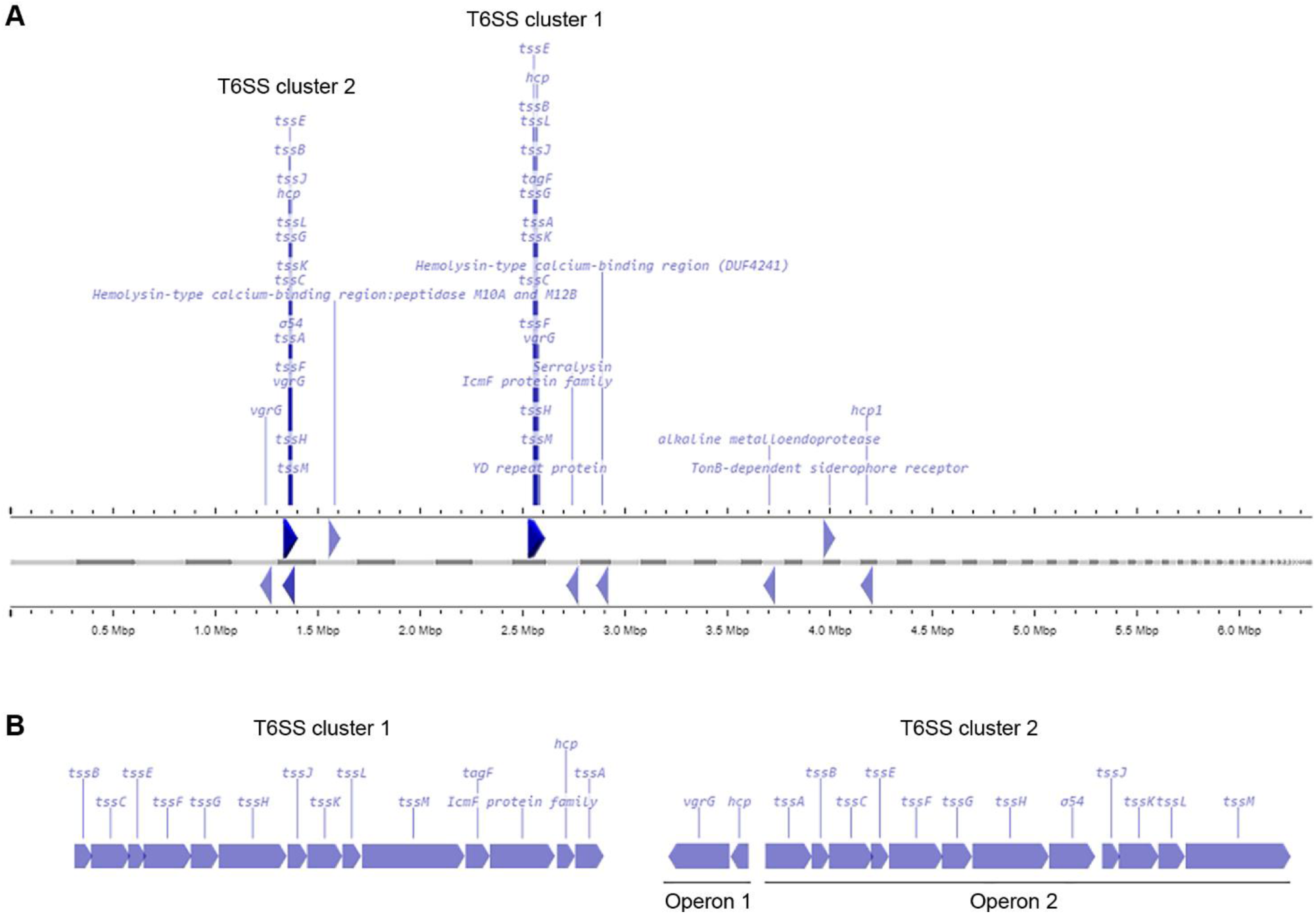
Genetic arrangement of T6SS genes in the genome of strain P21. (A) Arrangement of genes encoding for putative T6SS proteins in the genome of *P. syringae* pv. *aptata* strain P21. Gray bars indicate contigs ordered by length, ruler bars show the size of the genome. (B) Magnified genetic arrangements of the two T6SS clusters.

### *P. syringae* pv. *aptata* strains utilize distinct secretion-based infection strategies correlated with pathogenicity outcome

To get more in-depth insights into the patterns of protein secretion and identify potential new secreted proteins, we clustered all strains and the analyzed proteins, based on the difference of the intensity of the secreted proteins from the respective average value across all runs (Fig. 3). As already notable for many proteins in the previous comparisons (Table 2), the strains clustered in three clades: (i) the low virulence strains P16 and P17, (ii) the medium and high virulence strains P21 and P26 and (iii) the medium and high virulence strains P23 and P93.

**Figure 3.**
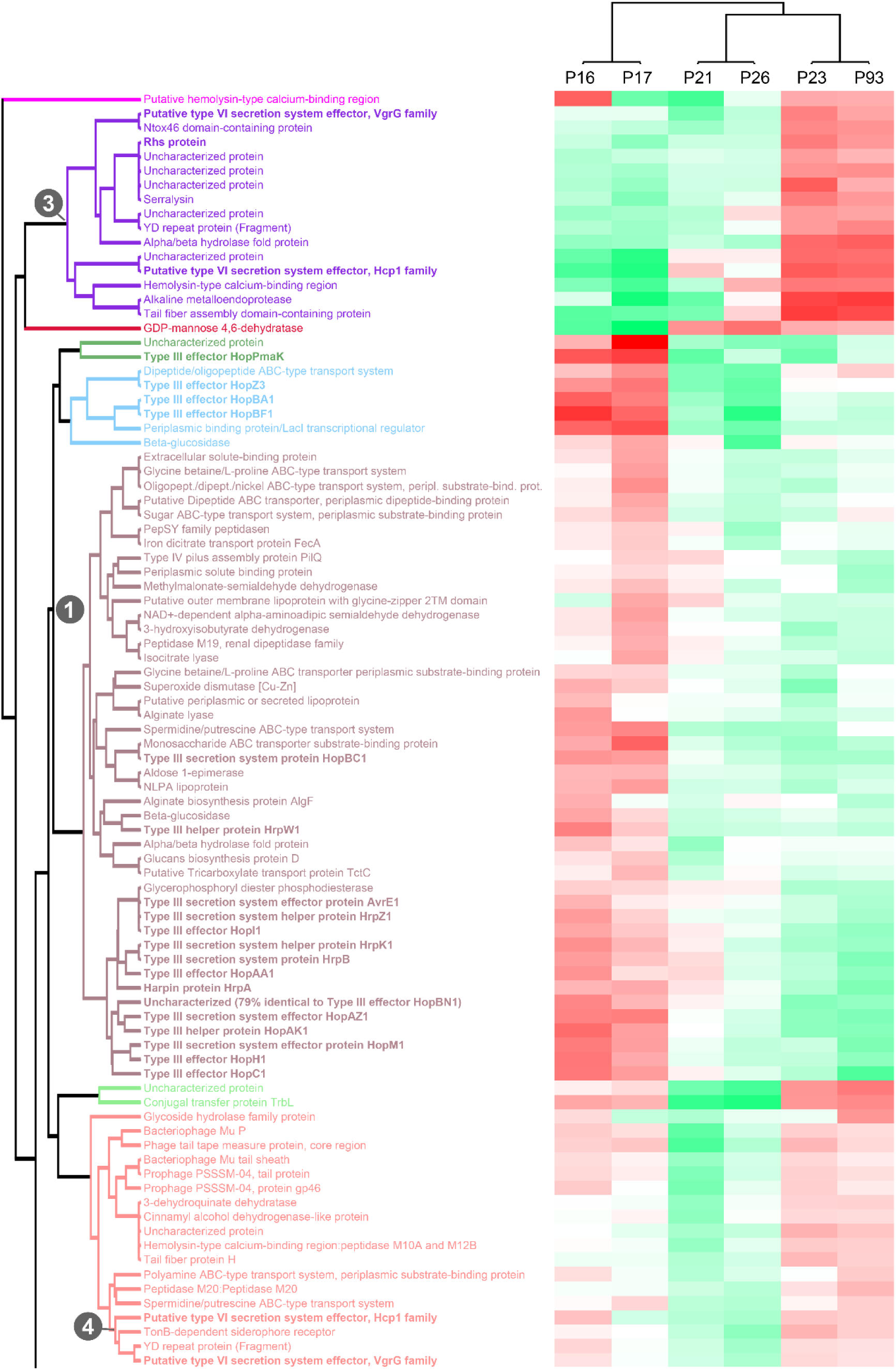
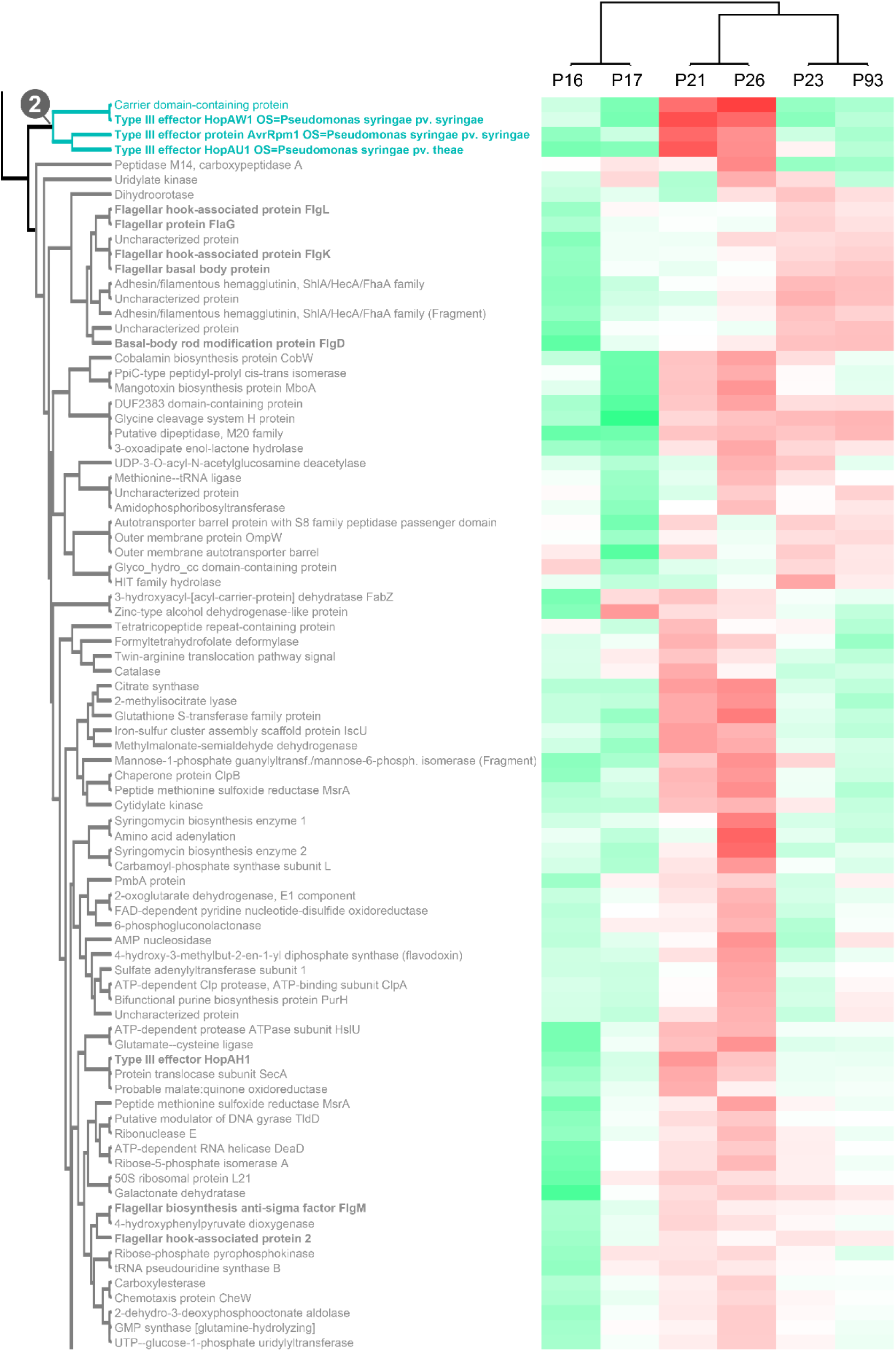
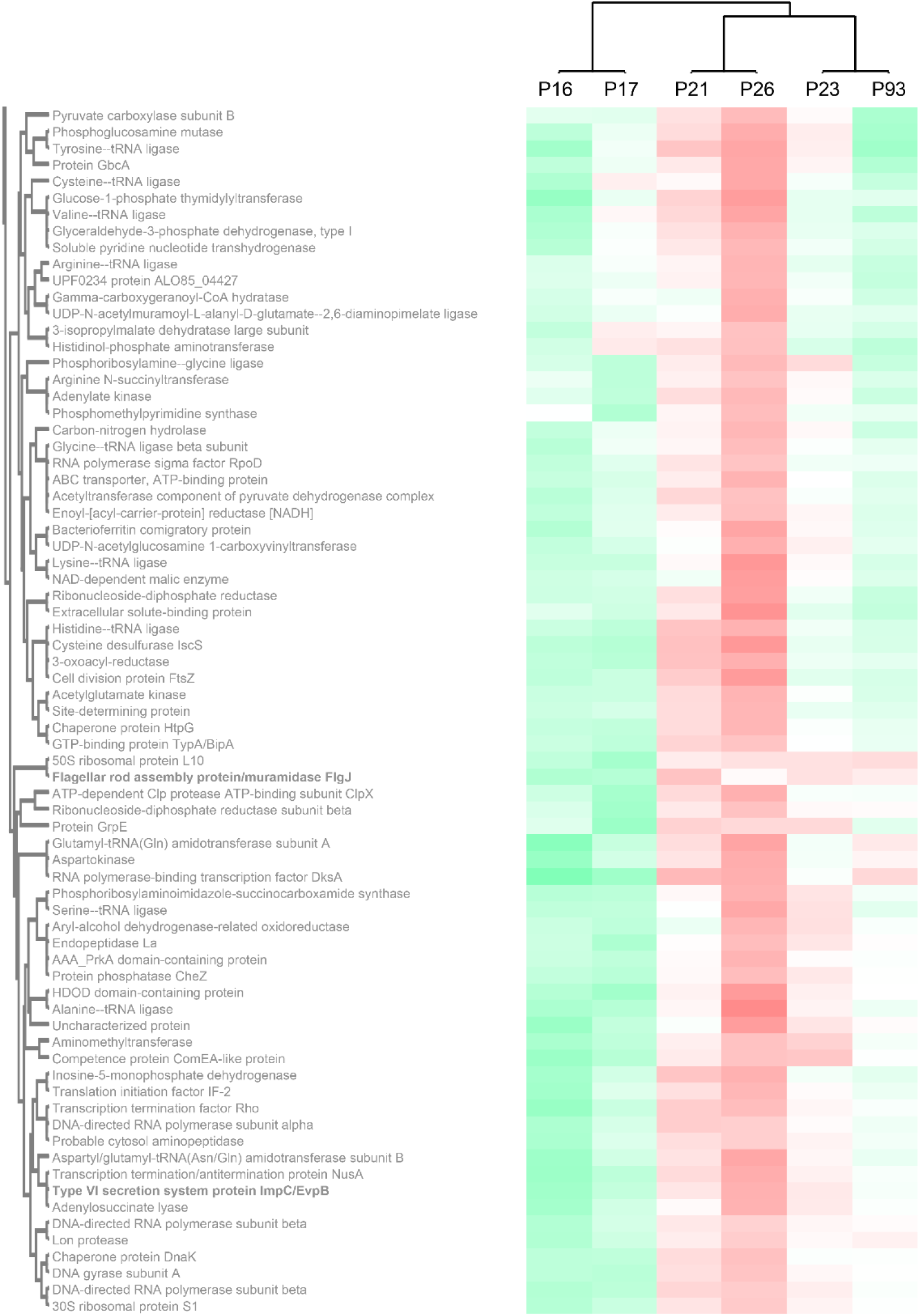
Clustering of strains and proteins in the culture supernatant indicates co-regulation of secretion of proteins in different strains. Clustering of all strains and secreted proteins previously analyzed with a maximal pairwise log 2 intensity difference of ≥3, based on the respective deviations from the average intensity. Color scale indicates average protein intensities. Different shades of red indicate increased protein intensity in the respective supernatant, whereas shades of green indicate lower intensities (median of 2 biological replicates). Strains and proteins were clustered in Perseus, using the following settings: Euclidean distance; average linkage; preprocess with k-means; number of clusters: 3 (columns), 150 (rows); 10 row clusters for coloration.

Flagellar exported proteins, in line with being more uniformly exported in all strains, fall in a large clade of proteins with limited difference between the strains.

In contrast, the clustering clearly displayed the co-regulation of secretion of the majority of the T3SS-secreted proteins (aubergine/green/blue clade (1) in Fig. 3) and a distinct cluster of T3SS effectors with different intensity distributions (turquoise clade (2) in Fig. 3), suggesting a different regulation of the secretion of these two subsets of T3SS effectors.

Similar to the T3SS export substrates, the known T6SS-secreted proteins cluster in two clades with distinct regulation (purple clade (3) and lower branch of pink clade (4) in Fig. 3, respectively; clade 4 includes the Hcp1 and VgrG proteins strongly secreted in all strains). Interestingly, each of these clades includes other proteins, whose presence in the culture supernatant follows a similar pattern between strains. Notably, some of these proteins are annotated as YD repeat proteins, which, like the related Rhs proteins, have been identified as T6SS effectors in other organisms (Koskiniemi et al., 2013; Pei et al., 2020), or as enzymes with a potential toxic activity that are candidates for T6SS effectors. To further explore this possibility, we screened the whole *P. syringae* pv. *aptata* proteome (5074 proteins) for potential T6SS effectors using the Bastion6 software (Wang et al., 2018) to predict T6SS effectors based on protein features, such as sequence profile (composition, permutation and combination modes of amino acids, orders of amino acids, similarities and homologies with other proteins), evolutionary information and physicochemical property. In line with our hypothesis, several of the proteins identified in our analysis are amongst the 251 proteins identified by Bastion6 as potential T6SS effectors (score value > 0.7), including several proteins from clades 3 and 4 (Suppl. Table 3). In particular, this was the case for the YD repeat protein (hit score 0.926), serralysin (0.791), and the alkaline metalloendoprotease (0.757) in clade 3, and the TonB-dependent siderophore receptor (0.791) and M10A/M12B hemolysin-type calcium-binding peptidase (0.742) in clade 4. All putative novel T6SS effectors are encoded separately from the T6SS gene clusters (Fig. 2), which is common among T6SS effectors (Barret et al., 2011). Additionally, we searched the genome of P21 for immunity protein-encoding genes in the vicinity of putative novel effectors and found several candidates. *Inh/omp19*, which encodes a predicted alkaline protease inhibitor, is located in a two-gene operon together with the gene encoding the alkaline metalloendoprotease. Similarly, Smi1, a homolog of the Tdi immunity protein (Zhang et al., 2011), is encoded in the neighboring operon of the gene encoding the YD repeat protein, in line with a role as an immunity protein. Taken together, these findings strongly support the hypothesis that some or all of these identified proteins are *bona fide* T6SS effectors.

## DISCUSSION

The establishment of a bacterial plant infection involves multiple virulence factors and corresponding host defense mechanisms, all of which are modulated by environmental factors (Morris et al., 2019). Understanding the pathogenic potential of *P. syringae* requires appreciating the structure, functionality, and mode of action of these virulence factors, especially at the crucial step when the pathogen enters into the plant apoplast environment. The main aim of this study was to evaluate the hypothesis that differences in pathogenicity between *P. syringae* pv. *aptata* strains correspond to variations in the repertoire and level of secreted virulence factors. To this aim, we performed functional assays and quantitative proteomics for six *P. syringae pv. aptata* strains with known pathogenic potential, focusing on proteins exported by the T3SS and T6SS, as well as flagellar proteins, which all are known to contribute to the pathogenic potential of bacterial plant pathogens (Pontes et al., 2020).

A recent genome-wide analysis in *P. syringae pv. aptata* P16 and P21 showed that these strains have 16 and 25 ORFs for T3SS effectors, respectively (Ranković et al., 2023), in line with a previous study finding 12 to 40 Hop/Avr open reading frames in various *P. syringae* pathovars (Baltrus et al., 2011). Here, using label-free quantification LC/MS-MS, we found that 23 different T3SS effectors were secreted into the extracellular fraction across the six tested *P. syringae* pv. *aptata* strains, confirming the secretion of most effectors detected in the genome-wide analysis (Ranković et al., 2023). Among the most abundant effectors were AvrE1 and HopM1, which have a crucial role in modulating humidity conditions after bacteria have entered the leaf apoplast (Xin et al., 2016), providing water-soaked spots, which were observed as an early disease symptom in previous *in planta* testing (Nikolić et al., 2018; Morris et al., 2019). Another highly abundant effector is HopBF1, involved in the phosphorylation of the molecular chaperone HSP90, which plays a role in pathogen-associated molecular pattern-triggered immunity, preventing activation of plant immunity response leading to the development of disease symptoms (Lopez et al., 2019).

The plant immune system can recognize the secretion of T3SS effectors by plant pathogens, which may result in effector-triggered immunity (ETI). As a response, plant pathogens have evolved mechanisms to avoid ETI, such as (i) mutations, recombination or loss of effector genes (Bundalovic-Torma et al., 2022), (ii) fine-tuning of secretion dosage of the core effectors (Laflamme et al., 2020), and (iii) metaeffector (effector-effector) interactions, where certain effectors suppress ETI triggered by other effectors (Martel et al., 2022). The low pathogenicity strains in our study (P16 and P17) show a significantly higher abundance of harpins, which could have a role in inducing a hypersensitive reaction (Choi et al., 2013) and well-known ETI-eliciting effectors (AvrE1, AvrRpm1, HopI1, HopZ3, HopAZ1 HopAA1 and HopBA1) (Laflamme et al., 2020), in comparison to high pathogenic strains. This suggests that the absence of fine-tuning in the secretion of core effectors (AvrE1, HopI1, HopAA1) and abundant secretion of effectors, which could lead to ETI, results in reduced pathogenicity. This hypothesis aligns with a recent finding that ETI in *Arabidopsis* was elicited upon overexpression of the *P. syringae* pv. *tomato* DC3000 effector AvrE1 (Laflamme et al., 2020). Furthermore, AvrE, HopI, HopZ, HopAZ, HopAA, and HopBA, which were abundantly secreted by the low pathogenicity strains P16 and P17 in our study, are members of effector families that elicit ETI in *Arabidopsis*.

Our study identified three T3SS effectors (AvrRpm1, HopAW1, and HopAU1) that were not detected in *P. syringae pv. aptata* so far, further broadening the effector spectrum of the pathovar. AvrRpm1 contributes to the suppression of plant defensive mechanisms by phosphorylation of the RIN4 protein, which represents a regulator of basal defense response in plants (Geng et al., 2016). HopAW1, also found in *P. syringae* pv. *phaseolicola*, is an effector protein containing a catalytic triad CHD motif, indicating cysteine protease activity (Mansfield, 2009). HopAU1 plays a role in the late stage of infection in *P. syringae* pv. *phaseolicola*, when it creates a replication-permissive niche in the apoplast, as Δ*hopAU1* strains display reduced growth in the plant apoplast (Macho et al., 2012). Quite suggestively, we found these effectors to be abundant in the secretome of strains P21 and P26, which have medium and high pathogenic features, indicating a possible role of these effectors in pathogenicity (aggressive virulence and broader host range) in a strain-specific manner.

In addition to T3SS-secreted effectors, we observed significant amounts of secreted T6SS components in the culture supernatant under apoplast-like conditions. T6SS in Gram-negative bacteria can be involved both in virulence and inter-bacterial competition (Gallegos-Monterrosa & Coulthurst, 2021; Unni et al., 2022). The number of T6SS clusters varies within the *P. syringae* species complex. While only one T6SS cluster is present in *P. syringae* pv. *actinidae, P. syringae* pv. *syringae* and *P. syringae* pv. *aesculi* (Bernal et al., 2018), the presence of two T6SS clusters was confirmed in *P. syringae* pv. *tomato, P. syringae* pv. *oryze* and *P. syringae* pv. *tabaci* (Sarris et al., 2010). Genomic mapping of strain P21 showed the existence of two different T6SS genetic islands in this genome (Fig. 2). The Hcp1 and VgrG proteins of cluster 1 were found to be secreted in high abundance by all strains, whereas another Hcp1, VgrG and a Rhs protein were distributed distinctly differently. By comparing the co-regulation of protein secretion in the tested strains, we identified several proteins as putative novel T6SS effectors, expanding the repertoire of putative effectors within the *P. syringae* species complex: Serralysin-like alkaline metalloproteases have also been detected to be co-regulated with T6SS proteins in *P. syringae* (Ali et al., 2022). The family of YD repeat proteins includes many bacterial toxins and proteins involved in cell-cell contact and communication, some of which are T6SS effectors (Koskiniemi et al., 2013). TonB-dependent siderophore receptors mediate substrate-specific transport across the outer membrane, mostly involved in the uptake of iron (Lin et al., 2017), in line with the recent finding that some T6SS effectors have a role in metal uptake when secreted into the extracellular environment and interact with TonB receptors allowing active transport of metal ions (Hernandez et al., 2020).

Our study shows a complex relationship between the function of T3SS and T6SS in *P. syringae pv. aptata* in apoplast-like conditions and suggests that the interplay between these two important systems shapes pathogenic strategies in a strain-specific manner. Strains P23 and P96 showed strong secretion of T6SS-related proteins and low secretion of most T3SS effectors, suggesting that in apoplast-like conditions, these strains rely more on the activity of the T6SS. On the other hand, the low-virulence strains P16 and P17 secreted higher amounts of most T3SS effectors, and lower amounts of T6SS-related proteins. Wang and colleagues found that T6SS mutants of *P. syringae* pv. *actinidae* had no pathogenicity phenotype *in planta*, but showed a strong down-regulation of several T3SS genes (Wang et al., 2021), highlighting the interplay of both secretion systems during infection. While further research is clearly needed to understand the role of the T6SS in *P. syringae* pathogenicity and its interaction with the T3SS, our results indicate that effector repertoire and secretion are correlated between the two secretion systems, contributing to distinct infection strategies.

Motility greatly impacts several aspects of *P. syringae* life cycle, especially in the transition from epiphytic to endophytic phase, and consequently to plant tissue invasion, pathogenicity, and disease symptoms development (Venieraki et al., 2016). In contrast to the results for T3SS- and T6SS-secreted proteins, we did not find a clear correlation between the abundance of flagellar proteins and swarming or swimming motility among tested strains, with the possible exception of strain P16, which showed low levels of exported flagellar components and the lowest motility (Suppl. Fig. 1).

This study was initially motivated by previous findings of large intra-pathovar diversity in pathogenicity within a strain collection. We reasoned that this divergence could be based on the differential secretion of effector proteins. Our secretome profiling of six *P. syringae pv. aptata* strains in apoplast-like conditions defines the T3SS effector repertoire and shows clear differences in the abundance of secreted effectors among strains with different pathogenic phenotypes. Based on the higher secretion level of conserved T3SS effectors in the low pathogenic strains, we hypothesize that *P. syringae* infection outcome depends on the fine-tuning of effector secretion, especially for ETI elicitors. In addition, we have expanded the spectrum of T3SS and T6SS effectors in *P. syringae* strains pathogenic on sugar beet, leading us to conclude that *P. syringae* pv. *aptata* may use protein secretion in a more diverse and flexible manner during infection than was previously known. Our analysis of the repertoire and abundance of secreted virulence factors between strains of different pathogenicity allows to distinguish three strategies in *P. syringae* strains pathogenic on sugar beet: (i) strong secretion of the majority of T3SS effectors/low secretion of T6SS effectors, (ii) strong secretion of differently regulated T3SS effectors/low secretion of T6SS effectors, and (iii) strong secretion of T6SS effectors/low secretion of all T3SS effectors. These patterns of secreted major virulence factors could contribute to a better understanding of overall virulence strategies of *P. syringae* and allow for tailored treatment of high-pathogenicity strains and pathovars of this important plant pathogen.

## Acknowledgements

Work in the research group of Andreas Diepold is supported by the Max Planck Society. This work was supported by the Ministry of Science, Technological Development and Innovations of Serbia, Grant No. 451-03-47/2023-01/200178 and Postdoc Fellowship No. 451-03-854/2019-14. The funders had no role in study design, data collection and analysis, decision to publish, or preparation of the manuscript. We gratefully acknowledge the help of Marco Herfurth, Max Planck Institute for Terrestrial Microbiology, Marburg, with electron microscopy imaging and of Daniel Unterweger, University of Kiel and Max Planck Institute for Evolutionary Biology, Plön, on the discussion of the T6SS data.

## Footnotes

1 Hrp - Hypersensitive reaction and pathogenicity

2 Avr - avirulence, Hop - Hrp-dependent outer protein

3 http://www.pseudomonas-syringae.org/; effectors were added from *P. syringae* pv. *syringae* B728a (UP000247706), *P. syringae* pv. *tomato* DC3000 (UP000002515), *P. syringae* pv. *theae* ICMP3934 (UP000282636), *P. syringae* pv. *phaseolicola* 1448A (UP000000551), *P. syringae* pv. *maculicola* ES4326 (UP000003811), *P. amygdaly* (formerly *P. syringae*) pv. *tabaci* ICMP2835 (UP000050478), *P. syringae* pv. *spinaciae* ICMP16929 (UP000050384), and *P. syringae* pv. *cilantro* 81035 (UP000037891)

