## Supplementary Information for "Repertoire and abundance of secreted virulence factors shape the pathogenic capacity of *Pseudomonas syringae* pv. *aptata*"

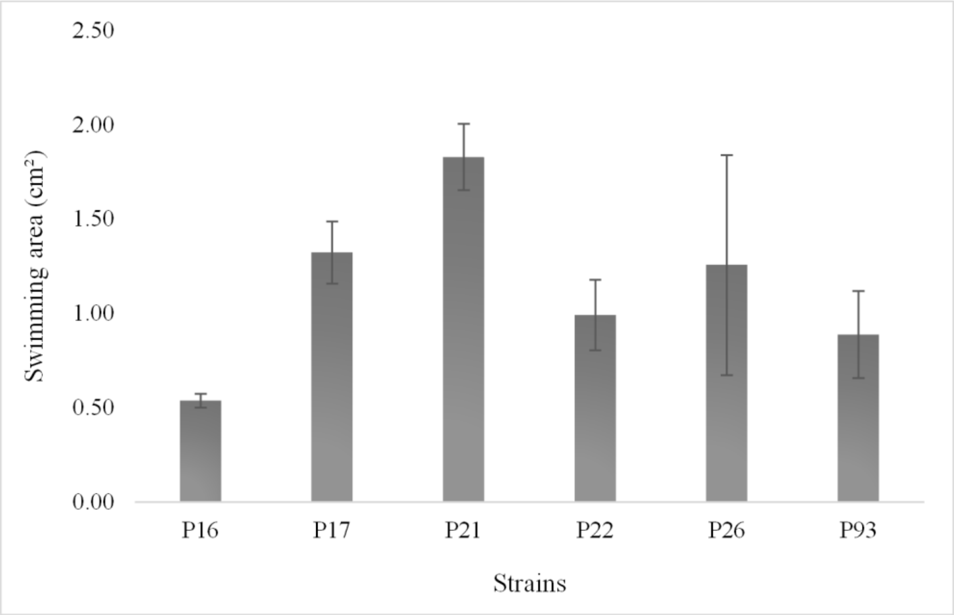


**Suppl. Fig. 1: Swimming areas of strains used in this study.**

Swimming diameters of *P. syringae* pv. *aptata* strain used in this study. $n = 3$; error bars denote standard deviation.

**Suppl. Table 1: Distribution of measured diameters of T3SS pili and flagella identified in electron microscopy experiments**

Diameters were measured manually using ImageJ (see material and methods for details), individual measurements are indicated.

| Identified as | Individual diameter (nm) | | | | | | | | | | Average diameter (nm) | St.dev. (nm) |
|---|---|---|---|---|---|---|---|---|---|---|---|---|
| **Pili** | 8.20 | 7.11 | 8.12 | 6.82 | 8.17 | 6.85 | 7.02 | 5.70 | 7.11 | 6.82 | **7.33** | 0.83 |
| **Flagella** | 23.86 | 17.20 | 16.29 | 14.76 | 18.35 | 16.62 | 12.56 | 25.68 | 15.69 | 19.12 | **18.01** | 4.02 |

**Suppl. Table 2: Additional information for Table 2, relative quantification of secreted proteins**

Measured intensities for proteins with at least five detected peptides and a maximal $log_2$ intensity difference between individual strains of ≤3. See Table 2 and main text for details.

| Protein | Average protein intensity (log 2) in indicated strains | | | | | | Overall avg. log 2 intens. | Max. diff. log 2 intens. |
|---|---|---|---|---|---|---|---|---|
| | P16 | P17 | P21 | P23 | P26 | P93 | | |
| **Type III secretion system** | | | | | | | | |
| HopAH2 protein | 25.226 | 26.209 | 25.993 | 25.101 | 26.396 | 24.214 | 25.523 | 2.182 |
| **Flagellum** | | | | | | | | |
| Flagellin | 34.499 | 35.046 | 35.588 | 36.593 | 35.820 | 35.922 | 35.578 | 2.094 |
| Flagellar hook protein FlgE | 29.056 | 29.485 | 29.506 | 31.566 | 30.591 | 31.278 | 30.247 | 2.511 |
| Flagellar basal-body rod protein FlgG | 26.871 | 27.631 | 27.577 | 29.343 | 28.457 | 29.406 | 28.214 | 2.536 |
| Flagellar hook-length control protein | 26.109 | 27.754 | 27.658 | 28.876 | 27.683 | 28.808 | 27.815 | 2.767 |

### Suppl. Table 3: List of potential T6SS effectors as determined by Bastion6

Bastion6 T6SS effector prediction results for proteins analyzed in proteomics analysis. Single method-based models and ensemble results, as indicated for all proteins with a prediction score larger than 0.7. T6SS-related proteins detected in the *P. syringae* pv. *aptata* secretome analysis are marked by bold font.

| Protein identifier | Protein annotation | Single Model Results | | | | | | | | | Ensemble Model Result Score |
|---|---|---|---|---|---|---|---|---|---|---|---|
| | | AAC | DPC | QSO | BLOSUM | DPC-PSSM | S-FPSSM | Pse-PSSM | CTDC | CTDT | |
| A0A0Q0C9C0 | Twin-arginine translocation pathway signal | 0.994 | 0.967 | 0.981 | 0.985 | 0.938 | 0.992 | 0.909 | 0.968 | 0.955 | 0.966 |
| A0A0Q0FP17 | Insecticidal toxin protein | 0.991 | 0.969 | 0.983 | 0.931 | 0.801 | 0.952 | 0.948 | 0.974 | 0.935 | 0.948 |
| A0A0Q0FP42 | DUF3274 domain-containing protein | 0.987 | 0.987 | 0.971 | 0.852 | 0.945 | 0.983 | 0.988 | 0.970 | 0.869 | 0.948 |
| A0A0Q0DDT3 | LPS-assembly protein LptD | 0.975 | 0.989 | 0.995 | 0.942 | 0.976 | 0.975 | 0.731 | 0.981 | 0.897 | 0.944 |
| A0A0Q0C8Q1 | CrtC domain-containing protein | 0.988 | 0.995 | 0.972 | 0.882 | 0.981 | 0.841 | 0.971 | 0.931 | 0.918 | 0.943 |
| A0A0Q0DLS3 | Insecticidal toxin protein | 0.992 | 0.965 | 0.984 | 0.855 | 0.987 | 0.838 | 0.972 | 0.948 | 0.897 | 0.939 |
| A0A0Q0C096 | Putative insecticidal toxin protein | 0.981 | 0.977 | 0.983 | 0.714 | 0.861 | 0.986 | 0.977 | 0.966 | 0.925 | 0.937 |
| A0A0Q0BX38 | Alginate biosynthesis protein AlgE | 0.969 | 0.987 | 0.995 | 0.935 | 0.963 | 0.966 | 0.876 | 0.885 | 0.902 | 0.937 |
| A0A0Q0CEY7 | Alginate lyase | 0.938 | 0.933 | 0.959 | 0.746 | 0.951 | 0.994 | 0.985 | 0.962 | 0.932 | 0.936 |
| A0A0Q0DLW9 | Insecticidal toxin protein | 0.976 | 0.953 | 0.956 | 0.919 | 0.844 | 0.980 | 0.968 | 0.946 | 0.892 | 0.936 |
| A0A0Q0DSL3 | Rhs family protein | 0.986 | 0.978 | 0.942 | 0.728 | 0.852 | 0.969 | 0.947 | 0.944 | 0.930 | 0.927 |
| A0A0Q0C688 | **YD repeat protein** | **0.994** | **0.921** | **0.969** | **0.601** | **0.870** | **0.916** | **0.992** | **0.977** | **0.966** | **0.926** |
| A0A0Q0CIG8 | Type III effector phosphothreonine lyase | 0.994 | 0.946 | 0.965 | 0.894 | 0.921 | 0.805 | 0.967 | 0.907 | 0.920 | 0.926 |
| A0A3M3EB84 | Type III effector HopAI1 | 0.993 | 0.941 | 0.942 | 0.885 | 0.919 | 0.839 | 0.964 | 0.904 | 0.926 | 0.925 |
| A0A0Q0D6F6 | Putative type VI secretion system effector, VgrG family | 0.953 | 0.946 | 0.893 | 0.904 | 0.926 | 0.973 | 0.967 | 0.936 | 0.866 | 0.925 |
| A0A0Q0D543 | Outer membrane porin OprE | 0.955 | 0.966 | 0.950 | 0.755 | 0.967 | 0.996 | 0.732 | 0.968 | 0.945 | 0.925 |
| A0A0Q0BXY6 | Outer membrane porin | 0.952 | 0.966 | 0.963 | 0.954 | 0.931 | 0.990 | 0.742 | 0.951 | 0.853 | 0.922 |
| A0A0Q0C5E5 | OprD family outer membrane porin | 0.976 | 0.985 | 0.984 | 0.981 | 0.924 | 0.993 | 0.648 | 0.917 | 0.869 | 0.920 |
| A0A0N8T848 | Glycoside hydrolase family 18 protein | 0.983 | 0.940 | 0.980 | 0.954 | 0.897 | 0.980 | 0.796 | 0.923 | 0.796 | 0.911 |
| A0A0Q0DBI7 | Outer membrane adhesin like proteiin | 0.943 | 0.993 | 0.949 | 0.790 | 0.904 | 0.916 | 0.788 | 0.934 | 0.905 | 0.910 |
| A0A0Q0DIJ7 | Anaerobically-induced outer membrane porin OprE | 0.908 | 0.981 | 0.980 | 0.977 | 0.954 | 0.997 | 0.694 | 0.811 | 0.918 | 0.909 |
| A0A0N8T849 | WW domain-containing protein | 0.992 | 0.996 | 0.998 | 0.944 | 0.983 | 0.269 | 0.925 | 0.958 | 0.946 | 0.909 |
| A0A0Q0DD34 | Catalase-peroxidase | 0.985 | 0.799 | 0.966 | 0.924 | 0.838 | 0.942 | 0.899 | 0.947 | 0.866 | 0.908 |
| A0A0N8T9E7 | Type VI secretion system effector, Hcp1 family | 0.961 | 0.909 | 0.905 | 0.871 | 0.969 | 0.867 | 0.989 | 0.876 | 0.871 | 0.908 |
| A0A0N8T980 | Zona occludens toxin | 0.951 | 0.967 | 0.933 | 0.853 | 0.938 | 0.786 | 0.792 | 0.921 | 0.933 | 0.907 |
| A0A0Q0DZB3 | Rhs protein | 0.939 | 0.928 | 0.941 | 0.804 | 0.940 | 0.917 | 0.907 | 0.961 | 0.815 | 0.905 |
| A0A0N8T7S5 | Type III effector | 0.978 | 0.971 | 0.981 | 0.769 | 0.957 | 0.850 | 0.955 | 0.829 | 0.885 | 0.905 |
| A0A0Q0FVM6 | RHS repeat-associated core domain-containing protein | 0.949 | 0.873 | 0.820 | 0.914 | 0.983 | 0.975 | 0.893 | 0.864 | 0.912 | 0.903 |
| A0A0Q0FN69 | Nucleoside-specific channel-forming protein Tsx | 0.914 | 0.994 | 0.978 | 0.670 | 0.888 | 0.921 | 0.904 | 0.879 | 0.916 | 0.902 |
| A0A0Q0DI86 | YD repeat-containing protein | 0.985 | 0.960 | 0.986 | 0.931 | 0.812 | 0.481 | 0.861 | 0.972 | 0.930 | 0.900 |

| | | | | | | | | | | | |
|---|---|---|---|---|---|---|---|---|---|---|---|
| A0A0Q0CV75 | Putative type VI secretion system effector, VgrG family | 0.857 | 0.913 | 0.905 | 0.945 | 0.922 | 0.945 | 0.896 | 0.909 | 0.840 | 0.898 |
| A0A0Q0C7Z5 | YD repeat-containing protein | 0.952 | 0.974 | 0.955 | 0.900 | 0.517 | 0.868 | 0.900 | 0.979 | 0.894 | 0.898 |
| A0A0Q0IAL8 | Porin | 0.994 | 0.991 | 0.987 | 0.815 | 0.913 | 0.969 | 0.649 | 0.970 | 0.756 | 0.897 |
| A0A0N8T9B6 | Type III helper protein HopAK1 | 0.868 | 0.989 | 0.984 | 0.976 | 0.952 | 0.992 | 0.915 | 0.653 | 0.904 | 0.895 |
| A0A0Q0FJN8 | Putative glycine-glutamate dipeptide porin OpdP | 0.947 | 0.960 | 0.987 | 0.612 | 0.919 | 0.989 | 0.769 | 0.946 | 0.836 | 0.893 |
| A0A0Q0D2R3 | YD repeat-containing protein | 0.991 | 0.985 | 0.993 | 0.941 | 0.816 | 0.427 | 0.832 | 0.972 | 0.895 | 0.892 |
| A0A0Q0E0H9 | Sucrose porin | 0.951 | 0.962 | 0.950 | 0.769 | 0.928 | 0.932 | 0.946 | 0.947 | 0.709 | 0.892 |
| A0A0Q0DL37 | Myo-inositol catabolism protein IolB | 0.987 | 0.974 | 0.904 | 0.675 | 0.866 | 0.838 | 0.810 | 0.931 | 0.918 | 0.892 |
| A0A0Q0BGR4 | Type III effector HopBB1 | 0.968 | 0.783 | 0.936 | 0.507 | 0.946 | 0.895 | 0.953 | 0.951 | 0.943 | 0.889 |
| A0A0Q0FHJ9 | Alkaline phosphatase | 0.985 | 0.953 | 0.953 | 0.394 | 0.822 | 0.978 | 0.856 | 0.911 | 0.920 | 0.881 |
| A0A0Q0C4L0 | TIGR03756 family integrating conjugative element prot. | 0.956 | 0.940 | 0.958 | 0.794 | 0.902 | 0.929 | 0.888 | 0.802 | 0.827 | 0.881 |
| A0A0N0GH35 | Type III effector HopF2 | 0.925 | 0.785 | 0.891 | 0.762 | 0.929 | 0.880 | 0.810 | 0.930 | 0.911 | 0.877 |
| A0A0Q0C296 | DUF1329 domain-containing protein | 0.934 | 0.938 | 0.938 | 0.850 | 0.933 | 0.818 | 0.814 | 0.869 | 0.813 | 0.877 |
| A0A0Q0CXH4 | phospholipase C | 0.982 | 0.874 | 0.983 | 0.652 | 0.788 | 0.995 | 0.782 | 0.980 | 0.774 | 0.876 |
| A0A0Q0IJ36 | DUF1329 domain-containing protein | 0.948 | 0.959 | 0.899 | 0.767 | 0.955 | 0.703 | 0.827 | 0.938 | 0.812 | 0.874 |
| A0A0Q0DAE5 | Outer membrane porin | 0.821 | 0.963 | 0.939 | 0.907 | 0.933 | 0.993 | 0.701 | 0.773 | 0.869 | 0.871 |
| A0A0Q0D6E9 | Pectate lyase/Amb allergen | 0.940 | 0.897 | 0.893 | 0.896 | 0.921 | 0.998 | 0.908 | 0.666 | 0.866 | 0.869 |
| A0A0Q0FUG0 | Putative 3-carboxymuconate cye | 0.918 | 0.934 | 0.947 | 0.794 | 0.602 | 0.748 | 0.846 | 0.962 | 0.893 | 0.869 |
| A0A0N8T9D8 | Outer membrane porin | 0.947 | 0.977 | 0.970 | 0.961 | 0.912 | 0.995 | 0.676 | 0.739 | 0.770 | 0.868 |
| A0A0Q0IAK6 | Lipoprotein | 0.984 | 0.906 | 0.814 | 0.594 | 0.916 | 0.819 | 0.955 | 0.837 | 0.902 | 0.864 |
| A0A0N8TA35 | 50S ribosomal protein L15 | 0.981 | 0.912 | 0.915 | 0.775 | 0.867 | 0.716 | 0.966 | 0.873 | 0.779 | 0.864 |
| A0A0Q0FW13 | Type III effector HopAH1 | 0.924 | 0.987 | 0.926 | 0.953 | 0.857 | 0.935 | 0.965 | 0.552 | 0.876 | 0.862 |
| A0A0Q0E0Z2 | Non-heme catalase KatN | 0.773 | 0.892 | 0.860 | 0.933 | 0.848 | 0.636 | 0.906 | 0.912 | 0.908 | 0.861 |
| A0A0Q0DNS3 | Outer membrane porin | 0.900 | 0.974 | 0.931 | 0.670 | 0.944 | 0.999 | 0.709 | 0.837 | 0.768 | 0.856 |
| A0A3M5WLB8 | Type III effector HopX1 | 0.967 | 0.837 | 0.893 | 0.886 | 0.900 | 0.763 | 0.719 | 0.840 | 0.863 | 0.856 |
| A0A0N8T9W4 | DUF2778 domain-containing protein | 0.948 | 0.871 | 0.761 | 0.660 | 0.842 | 0.960 | 0.917 | 0.902 | 0.816 | 0.855 |
| A0A0Q0IF84 | FlhE protein | 0.988 | 0.617 | 0.828 | 0.568 | 0.917 | 0.882 | 0.818 | 0.936 | 0.953 | 0.851 |
| A0A0Q0IFS3 | HopAH2 protein | 0.883 | 0.971 | 0.809 | 0.813 | 0.986 | 0.859 | 0.626 | 0.803 | 0.886 | 0.851 |
| A0A0Q0IT12 | Cupin domain-containing protein | 0.986 | 0.865 | 0.889 | 0.847 | 0.960 | 0.630 | 0.838 | 0.772 | 0.856 | 0.849 |
| A0A0Q0CG93 | Fimbrial protein | 0.968 | 0.763 | 0.901 | 0.860 | 0.950 | 0.845 | 0.991 | 0.844 | 0.675 | 0.849 |
| A0A0Q0C1R6 | Adhesin | 0.954 | 0.978 | 0.974 | 0.860 | 0.906 | 0.820 | 0.532 | 0.805 | 0.792 | 0.849 |
| A0A0Q0CVD4 | Sushi domain-containing protein | 0.600 | 0.856 | 0.868 | 0.812 | 0.864 | 0.817 | 0.817 | 0.974 | 0.914 | 0.849 |
| A0A0Q0C5H9 | Levansucrase LscA | 0.867 | 0.975 | 0.640 | 0.793 | 0.967 | 0.973 | 0.973 | 0.836 | 0.747 | 0.848 |
| A0A0N8T8T5 | Fimbrial bioproteinsis outer membrane usher protein | 0.937 | 0.973 | 0.870 | 0.754 | 0.844 | 0.841 | 0.583 | 0.869 | 0.854 | 0.848 |
| A0A0N8T7Y0 | Type III secretion system effector protein AvrE1 | 0.941 | 0.924 | 0.940 | 0.786 | 0.971 | 0.395 | 0.917 | 0.882 | 0.800 | 0.848 |
| A0A0Q0BXZ7 | TonB-dependent receptor | 0.986 | 0.997 | 0.992 | 0.967 | 0.481 | 0.891 | 0.232 | 0.957 | 0.843 | 0.845 |
| A0A0Q0BT79 | YD repeat protein | 0.985 | 0.778 | 0.739 | 0.797 | 0.786 | 0.973 | 0.986 | 0.921 | 0.698 | 0.843 |
| A0A0Q0BVQ5 | Hemolysin activator protein, HlyB family | 0.979 | 0.918 | 0.983 | 0.731 | 0.742 | 0.783 | 0.456 | 0.955 | 0.822 | 0.842 |
| A0A0Q0DT79 | Catalase | 0.945 | 0.940 | 0.908 | 0.675 | 0.663 | 0.682 | 0.554 | 0.970 | 0.927 | 0.841 |
| A0A0Q0FST6 | Glycoside hydrolase, family alpha amylase catal. subun. | 0.942 | 0.922 | 0.932 | 0.777 | 0.629 | 0.960 | 0.743 | 0.933 | 0.696 | 0.841 |
| A0A0Q0DST2 | Outer membrane porin | 0.880 | 0.929 | 0.919 | 0.652 | 0.951 | 0.991 | 0.700 | 0.807 | 0.771 | 0.840 |

| | | | | | | | | | | | |
|---|---|---|---|---|---|---|---|---|---|---|---|
| A0A0Q0IJ17 | Iron dicitrate transport protein FecA | 0.972 | 0.961 | 0.956 | 0.944 | 0.372 | 0.881 | 0.292 | 0.931 | 0.933 | 0.839 |
| A0A0Q0BVV7 | Super protein | 0.878 | 0.836 | 0.973 | 0.958 | 0.851 | 0.737 | 0.701 | 0.953 | 0.668 | 0.839 |
| A0A0Q0BT59 | DnaJ domain-containing protein | 0.821 | 0.819 | 0.758 | 0.932 | 0.798 | 0.953 | 0.922 | 0.757 | 0.869 | 0.838 |
| A0A0Q0CXV1 | Peptidase C13 superfamily protein | 0.978 | 0.990 | 0.939 | 0.812 | 0.279 | 0.942 | 0.601 | 0.873 | 0.895 | 0.837 |
| A0A0Q0DQM4 | Type VI secretion system effector, Hcp1 family | 0.924 | 0.660 | 0.815 | 0.812 | 0.953 | 0.930 | 0.993 | 0.793 | 0.785 | 0.837 |
| A0A0Q0BTQ0 | PA14 domain-containing protein | 0.908 | 0.987 | 0.919 | 0.538 | 0.972 | 0.728 | 0.714 | 0.870 | 0.790 | 0.836 |
| A0A0Q0CAC5 | Alginate lyase | 0.904 | 0.813 | 0.944 | 0.511 | 0.970 | 0.987 | 0.987 | 0.892 | 0.619 | 0.835 |
| A0A0Q0C453 | Deferrochelatase | 0.911 | 0.788 | 0.635 | 0.958 | 0.788 | 0.719 | 0.865 | 0.910 | 0.873 | 0.834 |
| A0A0Q0IMU7 | 4-phytase | 0.947 | 0.997 | 0.967 | 0.609 | 0.777 | 0.911 | 0.275 | 0.918 | 0.845 | 0.832 |
| A0A0Q0BX48 | Alginate biosynthesis protein AlgX | 0.971 | 0.837 | 0.921 | 0.626 | 0.911 | 0.451 | 0.756 | 0.941 | 0.861 | 0.832 |
| A0A0Q0C4H0 | Cytochrome c domain-containing protein | 0.959 | 0.876 | 0.882 | 0.610 | 0.847 | 0.730 | 0.882 | 0.879 | 0.760 | 0.831 |
| A0A0Q0D8S3 | Chitin-binding protein | 0.728 | 0.775 | 0.641 | 0.860 | 0.902 | 0.980 | 0.963 | 0.809 | 0.885 | 0.829 |
| A0A0Q0DEY0 | Copper resistance protein B | 0.940 | 0.983 | 0.963 | 0.709 | 0.907 | 0.535 | 0.775 | 0.737 | 0.845 | 0.828 |
| A0A0Q0FN00 | L-sorbosone dehydrogenase | 0.857 | 0.916 | 0.968 | 0.903 | 0.631 | 0.918 | 0.628 | 0.688 | 0.879 | 0.822 |
| A0A0Q0D919 | Tannase/feruloyl esterase family protein | 0.969 | 0.806 | 0.875 | 0.377 | 0.901 | 0.973 | 0.728 | 0.793 | 0.872 | 0.820 |
| A0A0Q0BYM2 | VCBS repeat-containing protein | 0.906 | 0.814 | 0.773 | 0.648 | 0.931 | 0.875 | 0.926 | 0.724 | 0.843 | 0.820 |
| A0A0Q0BWR0 | Autotransporting lipase, GDSL family protein | 0.961 | 0.992 | 0.970 | 0.761 | 0.844 | 0.776 | 0.317 | 0.785 | 0.826 | 0.818 |
| A0A0Q0D9W4 | Putative lipoprotein | 0.976 | 0.842 | 0.655 | 0.868 | 0.981 | 0.539 | 0.834 | 0.920 | 0.726 | 0.818 |
| A0A0N8T874 | Secreted protein | 0.921 | 0.880 | 0.859 | 0.673 | 0.941 | 0.856 | 0.706 | 0.659 | 0.884 | 0.817 |
| A0A0Q0IIE1 | Tail fiber protein H | 0.781 | 0.984 | 0.971 | 0.929 | 0.981 | 0.986 | 0.736 | 0.567 | 0.683 | 0.815 |
| A0A0Q0D451 | DUF4329 domain-containing protein | 0.957 | 0.878 | 0.750 | 0.743 | 0.632 | 0.564 | 0.783 | 0.893 | 0.913 | 0.815 |
| A0A1S6YAB7 | HopBD1 type III effector | 0.891 | 0.581 | 0.364 | 0.871 | 0.918 | 0.879 | 0.939 | 0.931 | 0.913 | 0.812 |
| A0A0Q0FVA2 | Aldose 1-epimerase | 0.938 | 0.967 | 0.977 | 0.585 | 0.571 | 0.443 | 0.688 | 0.952 | 0.853 | 0.812 |
| A0A0Q0ICK8 | Nucleoside-specific outer membrane channel protein Tsx | 0.702 | 0.945 | 0.900 | 0.365 | 0.771 | 0.933 | 0.762 | 0.899 | 0.842 | 0.809 |
| A0A0N8T952 | 30S ribosomal protein S19 | 0.912 | 0.832 | 0.913 | 0.219 | 0.863 | 0.742 | 0.909 | 0.857 | 0.861 | 0.809 |
| A0A0Q0DMX1 | ADP-ribosylating toxin | 0.831 | 0.872 | 0.845 | 0.807 | 0.944 | 0.876 | 0.773 | 0.664 | 0.790 | 0.809 |
| A0A0Q0BYJ4 | FGE-sulfatase domain-containing protein | 0.957 | 0.900 | 0.887 | 0.859 | 0.098 | 0.953 | 0.744 | 0.800 | 0.896 | 0.809 |
| A0A0Q0DGE1 | L-sorbosone dehydrogenase | 0.795 | 0.883 | 0.959 | 0.837 | 0.641 | 0.935 | 0.699 | 0.728 | 0.806 | 0.808 |
| A0A0Q0DHV7 | Neutral zinc metallopeptidase | 0.956 | 0.856 | 0.860 | 0.919 | 0.789 | 0.689 | 0.761 | 0.802 | 0.673 | 0.806 |
| A0A0Q0C079 | Superfamily protein | 0.864 | 0.633 | 0.954 | 0.962 | 0.856 | 0.702 | 0.658 | 0.897 | 0.715 | 0.806 |
| A0A0Q0IJF7 | Flagellar biosynthesis protein FlgJ | 0.852 | 0.822 | 0.890 | 0.370 | 0.829 | 0.847 | 0.763 | 0.806 | 0.907 | 0.805 |
| A0A0Q0BRB6 | Putative type VI secretion system effector, Hcp1 family | 0.899 | 0.892 | 0.920 | 0.896 | 0.965 | 0.932 | 0.995 | 0.707 | 0.419 | 0.805 |
| A0A0Q0BV33 | Outer membrane autotransporter barrel | 0.931 | 0.985 | 0.962 | 0.951 | 0.852 | 0.920 | 0.451 | 0.602 | 0.706 | 0.802 |
| A0A3M2XX23 | Type III secreted effector hopPmaA | 0.964 | 0.912 | 0.796 | 0.872 | 0.896 | 0.231 | 0.579 | 0.883 | 0.854 | 0.801 |
| A0A0Q0FGQ1 | Rhs family protein with PAAR motif | 0.812 | 0.825 | 0.850 | 0.676 | 0.239 | 0.758 | 0.852 | 0.934 | 0.942 | 0.799 |
| A0A0Q0IL40 | Homogentisate 1,2-dioxygenase | 0.932 | 0.916 | 0.576 | 0.795 | 0.817 | 0.734 | 0.653 | 0.812 | 0.838 | 0.794 |
| A0A0N8T7V9 | Type III secretion system helper protein HrpK1 | 0.826 | 0.866 | 0.820 | 0.520 | 0.972 | 0.840 | 0.791 | 0.752 | 0.771 | 0.793 |
| A0A0Q0FCB3 | Pili bioproteinsis outer membrane usher protein FimD | 0.772 | 0.928 | 0.941 | 0.837 | 0.860 | 0.941 | 0.528 | 0.739 | 0.662 | 0.791 |
| **A0A0Q0DMU9** | **TonB-dependent siderophore receptor** | **0.953** | **0.987** | **0.969** | **0.907** | **0.112** | **0.721** | **0.312** | **0.910** | **0.869** | **0.791** |
| **A0A0Q0CED4** | **Serralysin** | **0.809** | **0.978** | **0.962** | **0.830** | **0.892** | **0.927** | **0.463** | **0.580** | **0.778** | **0.791** |
| A0A0Q0DN93 | MltA-interacting MipA | 0.958 | 0.829 | 0.852 | 0.650 | 0.824 | 0.633 | 0.424 | 0.870 | 0.844 | 0.790 |

| | | | | | | | | | | | |
|---|---|---|---|---|---|---|---|---|---|---|---|
| A0A0Q0DMQ1 | Peptidoglycan hydrolase FlgJ | 0.707 | 0.723 | 0.689 | 0.421 | 0.911 | 0.834 | 0.897 | 0.866 | 0.925 | 0.789 |
| A0A0Q0C7K3 | Putative integral membrane protein | 0.702 | 0.971 | 0.872 | 0.854 | 0.804 | 0.992 | 0.586 | 0.570 | 0.846 | 0.789 |
| A0A0Q0C7G4 | RHS repeat-associated core domain-containing protein | 0.815 | 0.712 | 0.611 | 0.733 | 0.968 | 0.957 | 0.760 | 0.889 | 0.707 | 0.788 |
| Q4ZV88 | Type III effector HopAP1 | 0.890 | 0.851 | 0.704 | 0.719 | 0.512 | 0.773 | 0.943 | 0.819 | 0.800 | 0.787 |
| A0A0Q0C227 | RHS family protein | 0.948 | 0.756 | 0.556 | 0.774 | 0.728 | 0.968 | 0.985 | 0.760 | 0.728 | 0.787 |
| A0A0Q0E0I9 | Sucrose-6-phosphate hydrolase | 0.860 | 0.967 | 0.934 | 0.911 | 0.888 | 0.974 | 0.874 | 0.495 | 0.554 | 0.786 |
| A0A0Q0DGY4 | TonB-dependent siderophore receptor | 0.956 | 0.961 | 0.973 | 0.848 | 0.134 | 0.727 | 0.225 | 0.940 | 0.885 | 0.786 |
| A0A0N8T954 | Putative Type IV pilus-associated protein | 0.739 | 0.988 | 0.892 | 0.732 | 0.783 | 0.828 | 0.610 | 0.715 | 0.778 | 0.786 |
| A0A0Q0DXX0 | TonB-dependent siderophore receptor | 0.936 | 0.961 | 0.984 | 0.744 | 0.108 | 0.785 | 0.303 | 0.939 | 0.884 | 0.786 |
| A0A0Q0D5Y8 | TonB-dependent siderophore receptor | 0.954 | 0.994 | 0.994 | 0.701 | 0.123 | 0.817 | 0.285 | 0.900 | 0.870 | 0.782 |
| A0A0Q0FE60 | Penicillin amidase | 0.960 | 0.867 | 0.934 | 0.366 | 0.666 | 0.826 | 0.476 | 0.919 | 0.759 | 0.781 |
| A0A0Q0BV93 | Cysteine dioxygenase | 0.983 | 0.848 | 0.759 | 0.634 | 0.531 | 0.493 | 0.688 | 0.964 | 0.816 | 0.780 |
| A0A0N1JBR5 | Restriction endonuclease | 0.886 | 0.728 | 0.786 | 0.475 | 0.859 | 0.610 | 0.795 | 0.906 | 0.795 | 0.779 |
| A0A0Q0IKU0 | TonB-dependent siderophore receptor | 0.953 | 0.989 | 0.978 | 0.714 | 0.120 | 0.741 | 0.326 | 0.953 | 0.821 | 0.779 |
| A0A0Q0CY49 | Tat pathway signal:copper-resistance protein CopA | 0.542 | 0.866 | 0.779 | 0.916 | 0.643 | 0.835 | 0.663 | 0.853 | 0.817 | 0.776 |
| A0A0Q0FSL3 | TonB-dependent receptor | 0.783 | 0.979 | 0.960 | 0.575 | 0.734 | 0.960 | 0.289 | 0.769 | 0.795 | 0.776 |
| A0A0Q0DMX0 | Autotransp. barrel prot., S8 fam. Peptid. passenger dom. | 0.944 | 0.977 | 0.920 | 0.970 | 0.832 | 0.910 | 0.403 | 0.577 | 0.630 | 0.776 |
| A0A0Q0IC24 | Penicillin amidase | 0.960 | 0.618 | 0.944 | 0.306 | 0.647 | 0.859 | 0.561 | 0.926 | 0.861 | 0.776 |
| A0A0Q0FV00 | LPD7 domain-containing protein | 0.836 | 0.870 | 0.786 | 0.714 | 0.548 | 0.834 | 0.688 | 0.744 | 0.851 | 0.775 |
| A0A0Q0DPI0 | Ig-like_bact domain-containing protein | 0.693 | 0.885 | 0.919 | 0.708 | 0.953 | 0.928 | 0.534 | 0.603 | 0.818 | 0.774 |
| A0A0Q0BYH0 | BNR repeat-containing glycosyl hydrolase | 0.864 | 0.947 | 0.939 | 0.893 | 0.784 | 0.914 | 0.923 | 0.530 | 0.520 | 0.773 |
| A0A0Q0BTV2 | Oligopeptidase B | 0.975 | 0.754 | 0.958 | 0.644 | 0.473 | 0.963 | 0.396 | 0.787 | 0.819 | 0.772 |
| A0A0Q0DPI5 | Beta-glucosidase-related glycosyl hydrolase | 0.889 | 0.928 | 0.934 | 0.688 | 0.586 | 0.884 | 0.267 | 0.808 | 0.764 | 0.770 |
| A0A3M5WIF0 | Type III effector protein AvrB3 | 0.969 | 0.905 | 0.530 | 0.390 | 0.827 | 0.515 | 0.931 | 0.863 | 0.820 | 0.770 |
| A0A0Q0IH46 | PhcA | 0.824 | 0.655 | 0.864 | 0.599 | 0.761 | 0.374 | 0.918 | 0.907 | 0.819 | 0.769 |
| A0A0Q0FUW7 | Single-stranded DNA-binding protein | 0.964 | 0.957 | 0.944 | 0.770 | 0.331 | 0.348 | 0.434 | 0.850 | 0.906 | 0.768 |
| A0A0Q0DRB6 | Glucose dehydrogenase | 0.795 | 0.938 | 0.827 | 0.872 | 0.890 | 0.967 | 0.488 | 0.606 | 0.684 | 0.767 |
| A0A0Q0FSP4 | Ovule protein | 0.692 | 0.785 | 0.715 | 0.670 | 0.612 | 0.772 | 0.675 | 0.931 | 0.840 | 0.766 |
| A0A0Q0D665 | S-formylglutathione hydrolase | 0.970 | 0.940 | 0.748 | 0.364 | 0.611 | 0.799 | 0.419 | 0.785 | 0.937 | 0.765 |
| A0A0Q0DM93 | Esterified fatty acid cis/trans isomerase | 0.911 | 0.951 | 0.906 | 0.800 | 0.774 | 0.540 | 0.896 | 0.645 | 0.593 | 0.765 |
| A0A0Q0CVS2 | pectate lyase | 0.642 | 0.996 | 0.835 | 0.839 | 0.955 | 0.999 | 0.667 | 0.375 | 0.828 | 0.764 |
| A0A0Q0C504 | YD repeat-containing protein | 0.872 | 0.707 | 0.373 | 0.342 | 0.968 | 0.868 | 0.899 | 0.928 | 0.805 | 0.762 |
| A0A0Q0DS86 | PAAR domain-containing protein | 0.672 | 0.897 | 0.381 | 0.573 | 0.937 | 0.857 | 0.980 | 0.668 | 0.925 | 0.761 |
| A0A0N0X6U3 | Type III effector hopW1 | 0.935 | 0.870 | 0.614 | 0.923 | 0.932 | 0.120 | 0.475 | 0.857 | 0.863 | 0.760 |
| A0A0Q0DMN3 | TonB-dependent siderophore receptor | 0.865 | 0.993 | 0.953 | 0.818 | 0.133 | 0.687 | 0.220 | 0.858 | 0.899 | 0.760 |
| A0A0N8TA54 | Allergen V5/Tpx-1 related protein | 0.899 | 0.788 | 0.558 | 0.672 | 0.233 | 0.902 | 0.883 | 0.800 | 0.912 | 0.759 |
| A0A0N8T9G5 | 6-phosphogluconolactonase | 0.868 | 0.719 | 0.687 | 0.798 | 0.361 | 0.859 | 0.805 | 0.824 | 0.800 | 0.759 |
| A0A0Q0C4Y3 | TIGR03751 family conjugal transfer lipoprotein | 0.876 | 0.684 | 0.839 | 0.747 | 0.701 | 0.677 | 0.631 | 0.827 | 0.744 | 0.758 |
| A0A0Q0DMA8 | Quinoprotein glucose dehydrogenase | 0.899 | 0.918 | 0.914 | 0.907 | 0.836 | 0.987 | 0.476 | 0.626 | 0.497 | 0.758 |
| A0A0Q0DGE7 | Membrane protein involved in aromatic hydrocarbon degradation | 0.926 | 0.970 | 0.925 | 0.516 | 0.564 | 0.913 | 0.407 | 0.740 | 0.727 | 0.758 |
| A0A0Q0C4N1 | Replication protein A | 0.815 | 0.662 | 0.719 | 0.392 | 0.990 | 0.932 | 0.971 | 0.786 | 0.649 | 0.757 |

| | | | | | | | | | | | |
|---|---|---|---|---|---|---|---|---|---|---|---|
| A0A0N8T8P5 | **Alkaline metalloendoprotease** | **0.687** | **0.996** | **0.971** | **0.890** | **0.802** | **0.993** | **0.375** | **0.510** | **0.734** | **0.757** |
| A0A0Q0BZX0 | DUF3034 domain-containing protein | 0.959 | 0.506 | 0.893 | 0.643 | 0.757 | 0.961 | 0.446 | 0.774 | 0.785 | 0.756 |
| A0A0Q0CFT2 | Peptidase M14, carboxypeptidase A | 0.972 | 0.884 | 0.912 | 0.715 | 0.341 | 0.662 | 0.110 | 0.923 | 0.855 | 0.756 |
| A0A0Q0D672 | ZnMc domain-containing protein | 0.745 | 0.605 | 0.592 | 0.573 | 0.323 | 0.743 | 0.966 | 0.968 | 0.964 | 0.755 |
| A0A0Q0D4S0 | ABC transporter | 0.768 | 0.878 | 0.791 | 0.790 | 0.880 | 0.524 | 0.877 | 0.822 | 0.543 | 0.754 |
| A0A0N8T8I6 | Flagellar hook protein FlgE | 0.823 | 0.976 | 0.904 | 0.864 | 0.533 | 0.990 | 0.355 | 0.551 | 0.791 | 0.752 |
| A0A0Q0C264 | TonB system transport protein | 0.969 | 0.937 | 0.980 | 0.793 | 0.398 | 0.575 | 0.130 | 0.741 | 0.894 | 0.751 |
| A0A0Q0BSZ3 | Peptidase S1, chymotrypsin | 0.748 | 0.562 | 0.533 | 0.914 | 0.840 | 0.891 | 0.919 | 0.770 | 0.725 | 0.751 |
| A0A0Q0BWQ0 | Peptidase M23B | 0.927 | 0.859 | 0.862 | 0.742 | 0.394 | 0.128 | 0.434 | 0.963 | 0.920 | 0.750 |
| A0A0Q0FQV4 | Glycosyltransferase sugar-binding dom.-containing prot. | 0.794 | 0.769 | 0.869 | 0.406 | 0.893 | 0.947 | 0.894 | 0.696 | 0.613 | 0.750 |
| A0A0Q0FIC0 | Beta-glucosidase | 0.954 | 0.824 | 0.967 | 0.530 | 0.531 | 0.831 | 0.219 | 0.951 | 0.658 | 0.749 |
| A0A0N8T7T1 | Lipoprotein | 0.763 | 0.752 | 0.531 | 0.671 | 0.819 | 0.667 | 0.911 | 0.732 | 0.859 | 0.748 |
| A0A0N8TAA6 | Secreted protein | 0.945 | 0.615 | 0.914 | 0.421 | 0.876 | 0.618 | 0.884 | 0.935 | 0.499 | 0.747 |
| A0A0Q0DPF6 | Putative lipoprotein | 0.654 | 0.969 | 0.748 | 0.445 | 0.940 | 0.921 | 0.652 | 0.603 | 0.819 | 0.747 |
| A0A0Q0C1H9 | Rhs element Vgr protein | 0.630 | 0.650 | 0.733 | 0.860 | 0.825 | 0.908 | 0.806 | 0.676 | 0.763 | 0.747 |
| A0A0Q0FH11 | Alpha-2-macroglobulin | 0.841 | 0.846 | 0.904 | 0.849 | 0.945 | 0.521 | 0.497 | 0.748 | 0.603 | 0.747 |
| A0A0N8TA31 | Killer protein | 0.715 | 0.846 | 0.798 | 0.865 | 0.574 | 0.638 | 0.393 | 0.829 | 0.841 | 0.746 |
| A0A0Q0BZ89 | Beta-glucosidase | 0.861 | 0.848 | 0.859 | 0.684 | 0.560 | 0.851 | 0.236 | 0.893 | 0.708 | 0.746 |
| A0A0Q0BUZ1 | STN domain-containing protein | 0.947 | 0.980 | 0.933 | 0.778 | 0.112 | 0.709 | 0.305 | 0.847 | 0.755 | 0.743 |
| A0A0N8T920 | Secreted protein | 0.796 | 0.655 | 0.724 | 0.360 | 0.620 | 0.877 | 0.728 | 0.885 | 0.829 | 0.743 |
| A0A0Q0DMU2 | 3-oxo-C12-homoserine lactone acylase PvdQ | 0.942 | 0.849 | 0.829 | 0.750 | 0.563 | 0.875 | 0.623 | 0.665 | 0.643 | 0.743 |
| Q08I86 | Type III effector HopD | 0.640 | 0.896 | 0.713 | 0.604 | 0.949 | 0.822 | 0.623 | 0.565 | 0.886 | 0.742 |
| A0A0Q0C1J6 | **Hemolysin-type calcium-binding region:peptidase M10A and M12B** | **0.790** | **0.972** | **0.934** | **0.933** | **0.553** | **0.952** | **0.311** | **0.556** | **0.726** | **0.742** |
| A0A0Q0CEH2 | Outer membrane autotransporter barrel | 0.869 | 0.991 | 0.930 | 0.870 | 0.866 | 0.854 | 0.263 | 0.435 | 0.730 | 0.742 |
| A0A0N8T8D5 | Rhs family protein with PAAR motif | 0.708 | 0.734 | 0.433 | 0.439 | 0.784 | 0.951 | 0.991 | 0.798 | 0.822 | 0.742 |
| A0A0Q0ILE3 | Aldose-1-epimerase superfamily protein | 0.963 | 0.798 | 0.937 | 0.482 | 0.608 | 0.296 | 0.540 | 0.943 | 0.734 | 0.740 |
| A0A0Q0DM01 | Peptide chain release factor RF-3 | 0.610 | 0.917 | 0.799 | 0.691 | 0.836 | 0.829 | 0.662 | 0.717 | 0.661 | 0.740 |
| A0A0Q0ICD7 | Outer membrane ferripyoverdine receptor | 0.947 | 0.949 | 0.975 | 0.376 | 0.114 | 0.802 | 0.345 | 0.884 | 0.825 | 0.740 |
| A0A0Q0BR59 | Type III effector HopH1 | 0.717 | 0.778 | 0.701 | 0.489 | 0.778 | 0.856 | 0.986 | 0.778 | 0.636 | 0.739 |
| A0A0Q0IPQ3 | Outer membrane ligand receptor | 0.909 | 0.865 | 0.928 | 0.284 | 0.777 | 0.944 | 0.249 | 0.806 | 0.698 | 0.739 |
| A0A0Q0DNU1 | TonB-dependent outer membrane receptor | 0.928 | 0.982 | 0.978 | 0.784 | 0.115 | 0.665 | 0.181 | 0.920 | 0.716 | 0.739 |
| A0A0Q0DAM4 | DUF3828 domain-containing protein | 0.758 | 0.662 | 0.779 | 0.414 | 0.921 | 0.585 | 0.818 | 0.722 | 0.876 | 0.739 |
| A0A0Q0IK58 | Rhs family protein | 0.910 | 0.372 | 0.760 | 0.674 | 0.990 | 0.890 | 0.993 | 0.869 | 0.428 | 0.739 |
| A0A0Q0BYS1 | Phage capsid protein | 0.842 | 0.749 | 0.609 | 0.638 | 0.757 | 0.939 | 0.919 | 0.724 | 0.617 | 0.739 |
| A0A0N8TAD7 | Peptidase S9, prolyl oligopeptidase active site region | 0.977 | 0.878 | 0.960 | 0.673 | 0.595 | 0.571 | 0.517 | 0.594 | 0.779 | 0.738 |
| A0A0Q0BWB9 | Cbb3-type cytochrome c oxidase subunit | 0.972 | 0.955 | 0.960 | 0.902 | 0.330 | 0.596 | 0.867 | 0.428 | 0.726 | 0.738 |
| A0A0Q0DLM4 | Tol-Pal system protein TolB | 0.597 | 0.866 | 0.690 | 0.757 | 0.317 | 0.850 | 0.671 | 0.826 | 0.872 | 0.738 |
| A0A0Q0BRG7 | DUF4352 domain-containing protein | 0.746 | 0.476 | 0.625 | 0.809 | 0.859 | 0.564 | 0.648 | 0.898 | 0.856 | 0.737 |
| A0A0Q0DMZ9 | Abhydrolase_10 domain-containing protein | 0.736 | 0.685 | 0.790 | 0.867 | 0.813 | 0.450 | 0.502 | 0.767 | 0.862 | 0.737 |
| A0A0Q0C0M9 | DUF1989 domain-containing protein | 0.943 | 0.813 | 0.721 | 0.493 | 0.701 | 0.749 | 0.596 | 0.833 | 0.668 | 0.737 |
| A0A0Q0C4T2 | Alpha-1,4-glucan:maltose-1-phosphate maltosyltransferase | 0.810 | 0.822 | 0.865 | 0.709 | 0.436 | 0.777 | 0.431 | 0.852 | 0.723 | 0.736 |

| | | | | | | | | | | | |
|---|---|---|---|---|---|---|---|---|---|---|---|
| A0A0Q0D8X1 | DUF1852 domain-containing protein | 0.959 | 0.581 | 0.545 | 0.245 | 0.791 | 0.648 | 0.888 | 0.938 | 0.806 | 0.736 |
| A0A0Q0C7S9 | Putative type VI secretion system effector, VgrG family | 0.848 | 0.871 | 0.883 | 0.887 | 0.893 | 0.406 | 0.801 | 0.592 | 0.592 | 0.735 |
| A0A0Q0IQW5 | Syringomycin biosynthesis enzyme 2 | 0.941 | 0.765 | 0.902 | 0.452 | 0.379 | 0.006 | 0.795 | 0.939 | 0.908 | 0.734 |
| A0A0Q0D1M6 | TonB-dependent siderophore receptor | 0.845 | 0.969 | 0.908 | 0.818 | 0.139 | 0.651 | 0.259 | 0.816 | 0.833 | 0.733 |
| Q88BH0 | Type III effector HopK1 | 0.885 | 0.776 | 0.727 | 0.801 | 0.766 | 0.380 | 0.639 | 0.918 | 0.584 | 0.731 |
| A0A0Q0C6G4 | VirK family protein | 0.859 | 0.841 | 0.832 | 0.564 | 0.721 | 0.664 | 0.909 | 0.782 | 0.479 | 0.730 |
| A0A3M5X1E1 | Type III effector HopT1-1 | 0.907 | 0.592 | 0.353 | 0.433 | 0.848 | 0.565 | 0.813 | 0.921 | 0.890 | 0.729 |
| A0A0Q0DIF8 | Rhs family protein | 0.884 | 0.783 | 0.624 | 0.317 | 0.502 | 0.913 | 0.903 | 0.761 | 0.763 | 0.728 |
| A0A0Q0C763 | Prepilin | 0.748 | 0.958 | 0.751 | 0.667 | 0.767 | 0.934 | 0.690 | 0.498 | 0.702 | 0.728 |
| A0A0Q0FN94 | TonB-dependent siderophore receptor | 0.951 | 0.945 | 0.942 | 0.801 | 0.137 | 0.624 | 0.193 | 0.797 | 0.797 | 0.727 |
| A0A0Q0D786 | Aconitase B | 0.868 | 0.393 | 0.760 | 0.319 | 0.864 | 0.642 | 0.702 | 0.963 | 0.789 | 0.727 |
| A0A0Q0D8W4 | Putative signal peptide protein | 0.884 | 0.913 | 0.971 | 0.331 | 0.878 | 0.790 | 0.325 | 0.782 | 0.573 | 0.727 |
| A0A0Q0CVW3 | Type III effector HopZ3 | 0.873 | 0.767 | 0.735 | 0.901 | 0.528 | 0.305 | 0.683 | 0.733 | 0.821 | 0.724 |
| Q87Y16 | Type III effector protein AvrPto1 | 0.583 | 0.811 | 0.699 | 0.498 | 0.912 | 0.806 | 0.649 | 0.743 | 0.768 | 0.723 |
| A0A0Q0DXX3 | DNA/RNA non-specific endonuclease | 0.758 | 0.736 | 0.717 | 0.642 | 0.426 | 0.835 | 0.600 | 0.878 | 0.733 | 0.723 |
| A0A0N8TAB7 | Glucans biosynthesis protein D | 0.850 | 0.818 | 0.859 | 0.546 | 0.455 | 0.498 | 0.535 | 0.765 | 0.859 | 0.721 |
| A0A0Q0IP33 | DNA-binding transcriptional activator OsmE | 0.799 | 0.647 | 0.804 | 0.654 | 0.623 | 0.548 | 0.830 | 0.860 | 0.640 | 0.721 |
| A0A0Q0DLQ9 | N-acetylmuramoyl-L-alanine amidase | 0.914 | 0.815 | 0.773 | 0.523 | 0.494 | 0.795 | 0.950 | 0.681 | 0.595 | 0.721 |
| A0A0N8T9T9 | Putative secreted protein | 0.879 | 0.667 | 0.764 | 0.651 | 0.927 | 0.832 | 0.861 | 0.534 | 0.608 | 0.720 |
| A0A0Q0E0W9 | CigR | 0.506 | 0.809 | 0.825 | 0.460 | 0.712 | 0.925 | 0.804 | 0.584 | 0.858 | 0.720 |
| A0A0Q0DJZ6 | Sugar ABC-type transport system, periplasmic substrate-binding protein | 0.831 | 0.816 | 0.909 | 0.710 | 0.174 | 0.496 | 0.541 | 0.862 | 0.786 | 0.719 |
| A0A0Q0DSY5 | Putative periplasmic ligand-binding protein | 0.698 | 0.967 | 0.688 | 0.765 | 0.686 | 0.851 | 0.435 | 0.549 | 0.827 | 0.719 |
| A0A0Q0ID75 | Autotransporter barrel protein with phosphatase-like passenger domain | 0.719 | 0.962 | 0.903 | 0.828 | 0.828 | 0.800 | 0.380 | 0.555 | 0.618 | 0.719 |
| A0A0N8T8C2 | Glucans biosynthesis protein G | 0.723 | 0.741 | 0.811 | 0.875 | 0.572 | 0.498 | 0.553 | 0.731 | 0.802 | 0.716 |
| A0A2R3F5Q2 | Type III effector avrA1 | 0.849 | 0.662 | 0.696 | 0.440 | 0.810 | 0.589 | 0.677 | 0.854 | 0.716 | 0.716 |
| A0A0Q0FKW3 | Peptidase aspartic, active site protein | 0.839 | 0.825 | 0.976 | 0.820 | 0.101 | 0.418 | 0.355 | 0.941 | 0.751 | 0.716 |
| A0A0Q0FGW7 | Type III effector HopI1 | 0.832 | 0.732 | 0.724 | 0.718 | 0.337 | 0.596 | 0.726 | 0.731 | 0.845 | 0.715 |
| A0A0Q0ILZ0 | Imelysin, Metallo peptidase, MEROPS family M75 | 0.712 | 0.855 | 0.727 | 0.963 | 0.839 | 0.350 | 0.608 | 0.656 | 0.722 | 0.715 |
| A0A0Q0DIF4 | Oxidoreductase alpha | 0.944 | 0.830 | 0.771 | 0.877 | 0.086 | 0.509 | 0.062 | 0.934 | 0.887 | 0.714 |
| A0A0Q0DQV9 | Pirin | 0.694 | 0.812 | 0.477 | 0.833 | 0.812 | 0.704 | 0.377 | 0.891 | 0.706 | 0.714 |
| A0A0Q0DP26 | Phenol degradation meta-pathway protein | 0.884 | 0.789 | 0.849 | 0.808 | 0.856 | 0.799 | 0.234 | 0.774 | 0.474 | 0.713 |
| A0A0Q0C9U5 | DUF4105 domain-containing protein | 0.914 | 0.870 | 0.892 | 0.518 | 0.909 | 0.764 | 0.788 | 0.646 | 0.357 | 0.713 |
| A0A0Q0DAI4 | Conjugal transfer protein | 0.336 | 0.686 | 0.716 | 0.623 | 0.740 | 0.833 | 0.751 | 0.759 | 0.879 | 0.712 |
| A0A0Q0DMP2 | Putative Membrane protein | 0.822 | 0.853 | 0.891 | 0.536 | 0.939 | 0.968 | 0.932 | 0.400 | 0.464 | 0.711 |
| A0A0Q0BXL6 | RHS family protein | 0.821 | 0.713 | 0.772 | 0.662 | 0.725 | 0.984 | 0.936 | 0.465 | 0.602 | 0.710 |
| A0A0Q0DXG2 | Alpha-ketoglutarate-dependent dioxygenase AlkB | 0.916 | 0.516 | 0.813 | 0.594 | 0.483 | 0.478 | 0.707 | 0.860 | 0.763 | 0.709 |
| A0A0Q0C803 | Membrane-bound lytic murein transglycosylase A | 0.800 | 0.797 | 0.910 | 0.765 | 0.597 | 0.645 | 0.522 | 0.461 | 0.858 | 0.709 |
| A0A0Q0CVK2 | Lipoprotein | 0.538 | 0.603 | 0.606 | 0.663 | 0.757 | 0.821 | 0.719 | 0.726 | 0.882 | 0.709 |
| A0A0Q0FEQ9 | Thiamine pyrophosphate-binding protein | 0.940 | 0.710 | 0.713 | 0.454 | 0.901 | 0.767 | 0.783 | 0.592 | 0.630 | 0.708 |
| A0A0Q0DN10 | Transcriptional regulator | 0.867 | 0.538 | 0.682 | 0.491 | 0.591 | 0.652 | 0.584 | 0.856 | 0.842 | 0.708 |
| A0A0Q0DZ24 | Conjugal transfer protein | 0.821 | 0.660 | 0.867 | 0.805 | 0.308 | 0.227 | 0.238 | 0.933 | 0.961 | 0.708 |

| | | | | | | | | | | | |
|---|---|---|---|---|---|---|---|---|---|---|---|
| A0A0Q0DGR5 | Twin-arginine translocation pathway signal | 0.503 | 0.780 | 0.797 | 0.740 | 0.821 | 0.807 | 0.492 | 0.719 | 0.706 | 0.707 |
| A0A0Q0FI54 | GSDH domain-containing protein | 0.885 | 0.861 | 0.823 | 0.822 | 0.494 | 0.730 | 0.503 | 0.639 | 0.614 | 0.707 |
| A0A0Q0CIW2 | DUF4105 domain-containing protein | 0.916 | 0.777 | 0.833 | 0.264 | 0.870 | 0.752 | 0.785 | 0.615 | 0.602 | 0.706 |
| A0A0Q0BRH4 | ATPase | 0.833 | 0.402 | 0.577 | 0.800 | 0.935 | 0.832 | 0.834 | 0.638 | 0.693 | 0.706 |
| A0A0Q0BXK5 | Putative cytoplasmic protein | 0.931 | 0.860 | 0.684 | 0.680 | 0.631 | 0.457 | 0.340 | 0.653 | 0.871 | 0.705 |
| A0A0Q0FCL0 | SmpA/OmlA family outer membrane lipoprotein | 0.655 | 0.709 | 0.611 | 0.421 | 0.625 | 0.836 | 0.602 | 0.809 | 0.865 | 0.705 |
| A0A0Q0C4S4 | Translocation and assembly module subunit TamA | 0.972 | 0.841 | 0.894 | 0.426 | 0.443 | 0.539 | 0.188 | 0.925 | 0.704 | 0.705 |
| A0A0Q0IBZ8 | Allantoate amidinohydrolase | 0.867 | 0.715 | 0.682 | 0.677 | 0.748 | 0.345 | 0.582 | 0.903 | 0.636 | 0.704 |
| A0A0Q0DL55 | Putative lipoprotein | 0.721 | 0.854 | 0.859 | 0.111 | 0.366 | 0.826 | 0.391 | 0.842 | 0.909 | 0.704 |
| A0A0Q0DLX5 | Endolytic peptidoglycan transglycosylase RlpA | 0.760 | 0.954 | 0.804 | 0.357 | 0.315 | 0.554 | 0.227 | 0.872 | 0.939 | 0.703 |
| A0A0Q0C5C9 | FGE-sulfatase domain-containing protein | 0.821 | 0.795 | 0.576 | 0.552 | 0.824 | 0.986 | 0.926 | 0.562 | 0.545 | 0.702 |
| A0A0N8T802 | Protocatechuate 3,4-dioxygenase, alpha subunit | 0.740 | 0.691 | 0.565 | 0.694 | 0.662 | 0.427 | 0.329 | 0.909 | 0.911 | 0.701 |
| A0A0Q0FAW2 | Putative TraI family relaxase/helicase | 0.904 | 0.697 | 0.659 | 0.503 | 0.526 | 0.624 | 0.745 | 0.695 | 0.806 | 0.701 |
| A0A0Q0BVG8 | Peptide methionine sulfoxide reductase MsrA | 0.963 | 0.492 | 0.762 | 0.495 | 0.604 | 0.618 | 0.148 | 0.969 | 0.830 | 0.701 |
| A0A0Q0C2C2 | Cupin-like domain protein | 0.931 | 0.509 | 0.595 | 0.538 | 0.928 | 0.450 | 0.615 | 0.723 | 0.856 | 0.700 |
| A0A0N8T9X7 | Transporter | 0.731 | 0.770 | 0.705 | 0.499 | 0.801 | 0.857 | 0.638 | 0.591 | 0.741 | 0.700 |
